# Cortical determinants of loudness perception and auditory hypersensitivity

**DOI:** 10.1101/2024.05.30.596691

**Authors:** Kameron K Clayton, Matthew McGill, Bshara Awwad, Kamryn S Stecyk, Caroline Kremer, Desislava Skerleva, Divya P Narayanan, Jennifer Zhu, Kenneth E Hancock, Sharon G Kujawa, Elliott D Kozin, Daniel B Polley

**Affiliations:** Eaton-Peabody Laboratories, Massachusetts Eye and Ear, Boston MA 02114; Department of Otolaryngology – Head and Neck Surgery, Harvard Medical School, Boston MA 02114

**Keywords:** Autism, schizophrenia, hearing loss, aging, parvalbumin, inhibition stabilized network, hyperacusis, tinnitus, hallucination, gamma stimulation

## Abstract

Parvalbumin-expressing inhibitory neurons (PVNs) stabilize cortical network activity, generate gamma rhythms, and regulate experience-dependent plasticity. Here, we observed that activation or inactivation of PVNs functioned like a volume knob in the mouse auditory cortex (ACtx), turning neural and behavioral classification of sound level up or down over a 20dB range. PVN loudness adjustments were “sticky”, such that a single bout of 40Hz PVN stimulation sustainably suppressed ACtx sound responsiveness, potentiated feedforward inhibition, and behaviorally desensitized mice to loudness. Sensory sensitivity is a cardinal feature of autism, aging, and peripheral neuropathy, prompting us to ask whether PVN stimulation can persistently desensitize mice with ACtx hyperactivity, PVN hypofunction, and loudness hypersensitivity triggered by cochlear sensorineural damage. We found that a single 16-minute bout of 40Hz PVN stimulation session restored normal loudness perception for one week, showing that perceptual deficits triggered by irreversible peripheral injuries can be reversed through targeted cortical circuit interventions.

## Introduction

From the ticking of a watch to the roar of a waterfall, cochlear non-linear amplification compresses a million-million-fold range of signal intensity (120 dB) into a dynamic range that can be encoded by central auditory neurons.^1,2^ Where and how central neurons transition from representing acoustic sound level to the perceptual quality of loudness is unknown. One clue comes from whole cell and extracellular recordings in the auditory cortex (ACtx), where in contrast to the monotonic sound level response functions that predominate in the auditory nerve and subcortical nuclei, an increasingly large fraction of single units exhibit tuning to a narrow range of sound levels sculpted by the strength and timing of intracortical inhibition.^3–6^

Cortical representations of sound level are plastic. On millisecond timescales, ACtx neurons shift informative ranges of sound level encoding to match dynamic sound level statistics.^7^ One slower timescales of minutes to weeks, ACtx neurons shift preferred sound level encoding to match levels paired with direct stimulation of neuromodulatory nuclei^8^ or with behavioral reinforcement in auditory learning tasks.^9,10^ Thus, higher stages of central auditory processing leverage local inhibitory circuits to reformat the sound level code inherited from the periphery and brainstem, endowing sound level representations with plasticity while also potentially introducing vulnerabilities for abnormalities in loudness perception. Indeed, loudness is among the most fragile of all perceptual features where nearly 15% of adults have reduced sensitivity to soft sounds,^11^ while approximately 10% of adults report just the opposite problem – that moderate intensity sounds are experienced as uncomfortably loud and overwhelming.^12^ Loudness hypersensitivity (alternately described as loudness hyperacusis or sound sensitivity) in commonly reported in the context of aging or cochlear sensorineural hearing loss (SNHL), though it is also a cardinal feature of schizophrenia and various neurodevelopmental disorders including autism and Fragile X syndrome.^13,14^

We propose that ACtx parvalbumin-expressing GABAergic neurons (PVNs) play an essential role in the representation and plasticity of cortical sound level representations and the perception of loudness. PVNs likely supply the rapid inhibitory postsynaptic currents that dynamically shape sound level tuning and plasticity of individual ACtx neurons.^15–17^ Further, PVN hypofunction, destabilization, and disinhibition of recurrently connected networks of PVNs and excitatory pyramidal neurons has been identified as a root cause of sensory processing abnormalities in models of schizophrenia,^18,19^ autism,^20^ Fragile X syndrome,^21^ SNHL,^22,23^ and aging.^24^ Cortical PVNs are Janus-faced; as essential regulators of cortical plasticity, they are both a cause and solution for abnormal sensory processing that directly underlies perceptual disorders.^25^ Deterioration and disengagement of PVNs can disrupt inhibition-stabilized networks, producing cortical hyperactivity and promoting deleterious behaviors^26^ but their targeted stimulation may be able to rekindle inhibition and reverse cortical decline.^27,28^ Here, we demonstrate that a short period of 40Hz PVN stimulation is sufficient to suppress ACtx sound responses and desensitize mice with abnormal sensitivity to loudness, underscoring a critical role for ACtx PVNs in the encoding and perception of sound level.

## Results

### PVNs provide a volume knob for sound level representations in the auditory cortex

Ensembles of cortical neurons encode salient spectrotemporal features based on when spikes occur in time and which neurons are active.^29,30^ Spectrotemporal feature representations are high-dimensional, reflecting spiking dynamics arranged in several nested spatial and temporal scales.^31,32^ The overall magnitude of sound, the sound level, underlies all spectrotemporal features and provides a physical basis for a cardinal feature of sound perception – loudness. Here, we predicted that sound level is represented as a comparatively simple, low-dimensional “meter” that can be decoded without resolving which neurons are active, nor when spiking occurs (**Figure 1A**).

**Figure 1.**
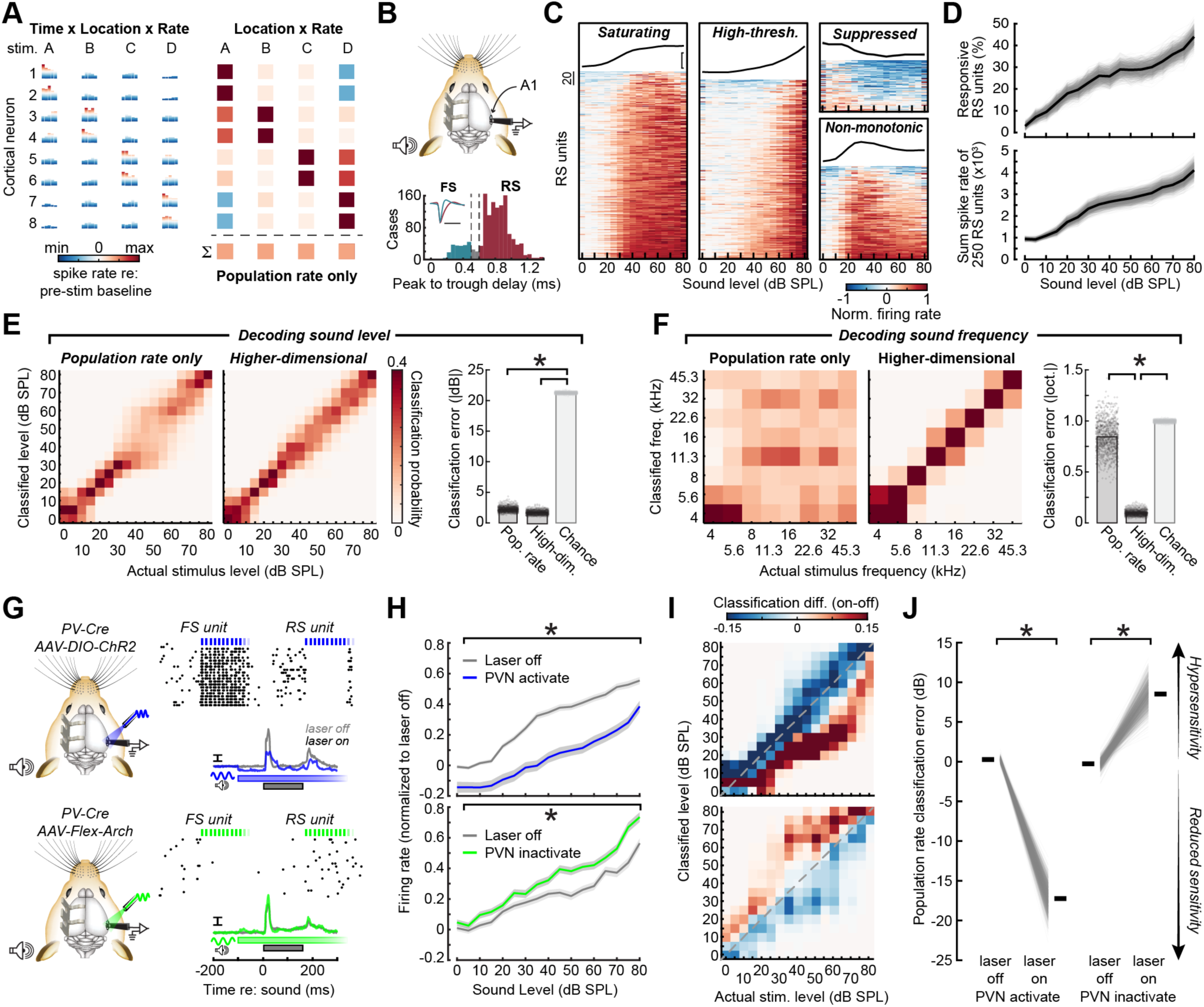
A low-dimensional population rate code for sound level in the primary auditory cortex. A) Cartoon data schematizes the spiking activity of 8 neurons in response to four stimuli. Stimulus representations can be high-dimensional, reflecting the location, timing, and rate of active neurons (*left*), the location and rate of active neurons (*right*), or a low-dimensional population rate code, reflecting only the summed rate of all neurons (*bottom-right*). B) A cartoon illustrates approach for extracellular recordings from A1 of unanesthetized, head-fixed mice with 64-channel silicon probes. Single unit waveforms were classified as fast-spiking (FS, teal) or regular spiking (RS, maroon) based on a bimodal distribution of peak to trough delays. C) Neurograms depict normalized, baseline-corrected spike rates for 721 sound-responsive RS units from 8 mice in response to broadband noise bursts of varying level. K-means clustering identified four distinct RS response profiles that can be described as saturating (n = 264), high-threshold (n = 252), suppressed (n = 74) and non-monotonic (n = 131). D) Percentage of significantly responsive RS units (*top)* and summed spike rate (*bottom*) for a random sample of 250 RS units as a function of increasing sound level. Thin lines represent each selection of 250 units. Thick line represents mean of 1000 selections. E) Confusion matrices depict decoding accuracy for sound level based solely on the population activity rate (*left*) versus a higher-dimensional neural representation (*middle*). Absolute value of sound level classification error based on summed activity rate or the higher-dimensional representation relative to the classification error that occurs by chance, with shuffled sound level labels. Decoding is performed on 1000 randomly drawn samples of 250 RS units (bar = mean). Asterisk indicates the conjunction of a p value < 0.05 after Bonferroni-Holm correction for multiple comparisons and an effect size (Hedge’s G) > 2. F) As per *E*, but applied to A1 RS unit decoding of tone pip frequency. G) Cartoon illustrates approach for documenting sound level processing in A1 RS units while PVNs are activated or inactivated with, ChR2 or Arch, respectively. Raster plots from four example units show a pair of directly activated and inactivated FS units (*top and bottom*, respectively) alongside synaptically suppressed (*top*) or activated (*bottom*) neighboring RS units. Laser pulses are 25ms in duration presented at 20Hz over a 0.5s period that subsequently tapered in amplitude to avoid rebound excitation. Mean RS PSTHs show sound-evoked spike rates with laser on and off. Scale bar = 2 sp/s. H) *Top*: Mean ±SEM spike rate-level functions are significantly suppressed by PVN activation (N/n = 4/401 mice/RS units). A 2x2 factorial linear mixed effects model ANOVA identified a significant main effect for PVN activation and a significant level x PVN activation interaction (F > 35, p < 3x 10^-9^ for each). *Bottom:* Steeper sound level growth with PVN inactivation (N/n = 4/320 RS units). A 2x2 factorial linear mixed effects model ANOVA identified a trend for PVN inactivation (F = 3.67, p = 0.06) but a significant level x PVN inactivation interaction term (F > 35, p < 2 x 10^-9^). I) Confusion matrices depict the change in classification probability between the laser off and laser on conditions based on the summed population activity rate. Other plotting conventions as per *E* and *F*. J) Classified sound level from summed population rate was nearly 20dB lower than actual sound level during PVN activation but nearly 10dB higher than the actual sound level during PVN inactivation. Decoding is performed on 1000 randomly drawn samples of 250 RS units for each condition (bar = mean). Asterisks indicate significant differences between laser on and off conditions based on a permutation test (p = 0.0001 for both).

To test this hypothesis, we made extracellular recordings of single units in the primary auditory cortex (A1) of head-fixed unanesthetized mice. Isolated units were classified as fast-spiking (FS) putative PVNs or regular spiking (RS) putative pyramidal neurons based on waveform shape (**Figure 1B**). We presented white noise bursts of varying intensity and observed substantial heterogeneity in sound level encoding at the level of individual RS neurons. Some neurons exhibited low-threshold saturating responses, others increased monotonically across a range of higher sound levels, were suppressed at intermediate and high levels, or were non-monotonic, resulting in tuning to a narrow range of sound levels (**Figure 1C-D**). Importantly, heterogeneity in single unit responses summed together to produce a linear population representation of sound level when measured either as the growth in the percentage of significantly responsive RS units (**Figure 1E**) or summed spike rate across randomly drawn samples of 250 RS units (**Figure 1F**).^9,33^

We found that sound level could be decoded using a PSTH-based classifier with < 3dB of error based on the summed population activity rate that disregarded spike timing and neuron identity (**Figure 1G, left**). Increasing the dimensionality of the level representation by allowing principal components related to unit identity into the decoding model offered minimal improvement in sound level identification (**Figure 1G, right**, statistical analyses here and elsewhere are reported in figure legends). By contrast, decoding a rudimentary spectrotemporal feature – the frequency of a pure tone – from the summed population activity rate was near chance and required the higher-dimensional representation to achieve accuracy (**Figure 1H**).

A representational scheme based solely on population activity rate would be robust micro-scale dynamics like representational drift.^34^ On the other hand, encoding strategies based on total network activity would be susceptible to changes in inhibitory tone that impose linear gain adjustments on cortical output.^35,36^ PVNs, in particular, impose bi-directional linear gain control on excitatory neurons,^37,38^ suggesting that dips or surges in their activity state could bias cortical level representations towards over-or under-reporting the physical sound level, respectively. To test this hypothesis, we optogenetically activated or inactivated PVNs with ChR2 or Archaerhodopsin-3 (Arch) to suppress or increase the excitability of RS units, respectively (**Figure 1I**). We observed that linear sound level representations in RS units were significantly suppressed by PVNs activation and significantly elevated during PVNs inactivation (**Figure 1J**). By training the RS unit population activity rate decoder on laser-off trials and then decoding the sound level from individual trials with PVNs activated or inactivated, we observed a bi-directional shift in sound level classification (**Figure 1K**). PVN activation caused sounds to be classified as 17.25 dB lower than their actual level, while PVN inactivation introduced an 8.52 dB bias in the opposite direction (**Figure 1L**).

It is worth noting that the recurrent connectivity of PVNs can produce effects consistent with inhibition stabilized networks, wherein PVN inactivation can cause excitation of neighboring PVNs and PVN activation can cause inhibition of neighboring PVNs.^39,40^ These paradoxical effects are prominent in preparations that activate large fraction of neurons at longer durations.^41,42^ By relying on shorter pulses of light at moderate laser powers in mice with restricted virus-mediated transduction of PVNs, we were able to manipulate local networks of PVNs and excitatory neurons in their linear operating rate without recruiting inhibition stabilization as the dominant motif.

### PVN activation – not sensory stimulation – at 40Hz elicits an endogenous gamma rhythm based on synchronization of A1 RS and FS unit spiking

In some circumstances, direct activation of PVNs can induce a sustained change in local circuit processing and cognition that outlasts the period of stimulation.^43,44^ PVN stimulation at 40-60Hz has proven particularly effective in eliciting robust gamma rhythms and enhancing interareal coupling for a period of 24 hours after stimulation is discontinued.^28^ Some studies have suggested that sensory stimulation can substitute for direct neurostimulation by producing gamma rhythms and imparting sustained changes in cortical microcircuits that can forestall or even reverse neurodegeneration and improve cognition,^27^ though the generality of these claims are a matter of debate.^45^

As a next step, we investigated whether these concepts could be applied to the neural representation of sound level in A1. Auditory thalamic neurons can reliably synchronize to 40Hz modulations in the sound pressure envelope, though most cortical neurons cannot reliably synchronize to 40 Hz stimulus modulations and instead utilize non-synchronized rate coding to encode higher ranges of modulation rates.^46^ Because local field potential (LFP) recordings reflect a strong presynaptic contribution, we predicted that presentation of sounds that flutter at 40Hz would entrain the LFP measured in A1 but not the endogenous RS and FS units. Instead, we predicted that strong endogenous recruitment of RS and FS spiking at gamma frequencies would require direct activation of PVNs at gamma frequencies.

We tested these predictions by measuring the LFP power spectrum and spiking activity of RS and FS units from all layers of A1 in response to 40 Hz acoustic stimulation, constant optogenetic PVN activation, and 40 Hz optogenetic PVN activation. We found that 40Hz acoustic stimulation elicited an intense band of A1 LFP gamma activity (**Figure 2B, top**). However, the spike trains of underlying FS units synchronized to the 40Hz sound whereas RS units exhibited a mixture of synchronized and non-synchronized spiking that resulted in increased power across multiple frequency bands (**Figure 2B, middle and bottom**). One second of continuous PVN activation strongly activated PVNs and suppressed RS unit spiking but did not elicit any structured gamma activity in the LFP or single unit spiking (**Figure 2C**). Activating PVNs at 40 Hz activated PVNs and suppressed RS units but also modulated their spike rates at 40 Hz, producing a selective gamma band increase in both the LFP and spiking power spectrum (**Figure 2D**). We quantified gamma entrainment as the power at the 30-50Hz gamma range relative to other frequencies withing 0-70Hz range and confirmed that both 40 Hz acoustic and 40 Hz PVN activation elicited significant LFP gamma entrainment (**Figure 2E**). However, RS unit entrainment with 40 Hz PVN activation was significantly greater than acoustic stimulation, which was no different than steady PVN activation (**Figure 2F**). These findings demonstrate that 40Hz PVN stimulation recruits strong entrainment of endogenous spiking in A1 RS units while acoustic stimulation at 40 Hz does not (**Figure 2E-F**).^46^

**Figure 2.**
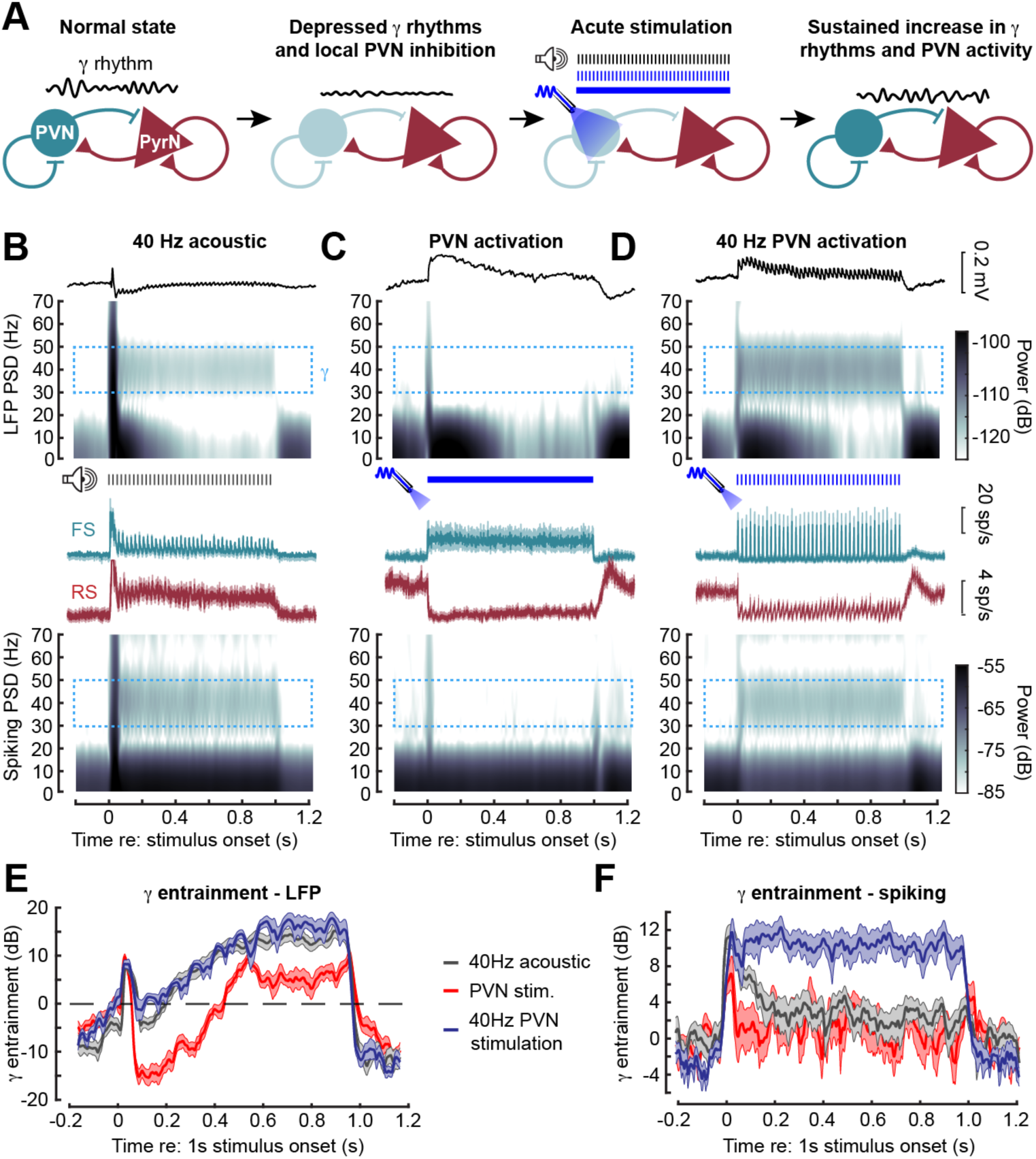
PVN activation – not sensory stimulation – entrains endogenous spiking at gamma frequencies. A) Cartoon illustrates prominent gamma rhythms associated with recurrently connected inhibitory PVNs and excitatory pyramidal neurons (PyrN). Diminished PVN activity is associated with reduced gamma rhythms, which could be reinvigorated and sustained through PVN activation at gamma frequencies. B) A1 local field potential (LFP) (top) and spiking activity (bottom) across layers during 40 Hz acoustic stimulation, Middle subpanel shows mean ±SEM FS and RS rates for 3 mice (58 FS units, 318 RS units). C) As per *B,* but with sustained PVN optogenetic activation D) As per *B*, but with 40 Hz PVN stimulation. E) LFP gamma entrainment was calculated as the PSD amplitude in the 35-45Hz range during the 1s stimulation period relative to other frequencies. LFP gamma entrainment with 40Hz acoustic and 40Hz PVN activation were significantly greater than PVN activation and not different from one another (N = 3 mice, 14 columnar recording sites; 2x2 repeated measures ANOVA main effect for time [F = 119.49, p = 7 x 10^-8^]; main effect for stimulus type [F = 30.63, p = 2 x 10^-7^]; time x stimulus type interaction [F = 21.9, p = 3 x 10^-6^]. Post-hoc pairwise contrasts: 40 Hz acoustic vs PVN (p = 5 x 10^-6^), 40Hz PVN vs PVN (p = 6 x 10^-5^), 40 Hz acoustic vs 40 Hz PVN (p = 0.44). F) As per *E* but measured from A1 spiking activity. Gamma entrainment to 40Hz PVN activation was significantly greater than PVN activation and 40 Hz acoustic activation, which were not different from each other (N = 3 mice, 14 penetrations, n = RS 376 units; 2x2 repeated measures ANOVA main effect for time [F = 35.43, p = 5 x 10^-5^]; main effect for stimulus type [F = 10.66, p = 5 x 10^-4^]; time x stimulus type interaction [F = 11.73, p = 3 x 10^-4^]. Post-hoc pairwise contrasts: 40 Hz acoustic vs PVN (p = 0.43), 40Hz PVN vs PVN (p = 0.002), 40 Hz acoustic vs 40 Hz PVN (p = 6 x 10^-4^).

### Sustained dampening of sound-evoked cortical activity with PVN gamma stimulation

Neurostimulation therapies are predicated on the idea that sustained changes in local circuit processing, perception, and cognition can arise from brief periods of structured neurostimulation.^47^ While interleaved PVN activation only shifted neural decoding of sound level towards reduced sensitivity while the laser was on (Figure 1J), we next asked whether inducing a sustained period of intense endogenous gamma activity would sustainably potentiate the influence of PVNs on local circuit processing after stimulation was discontinued. In this scenario, we hypothesized that sustained PVN potentiation would be associated with the following set of changes that we could measure with extracellular recordings: 1) enhanced LFP resting gamma power, 2) suppressed sound-evoked firing rates in RS units, 3) enhanced sound-evoked firing rates in FS units, and 4) enhanced PVN-mediated inhibition of spontaneous RS spike rates. We addressed these predictions by either activating PVNs with 1ms pulses of light at 40 Hz for 1000 seconds (PVN γ stimulation) or with a control protocol designed to phasically activate ACtx PVNs without inducing gamma activity (50ms pulse of light once per second for 1000 seconds, PVN control stimulation; **Figure 3A**). After a set of baseline measures and the 16-minute period of PVN γ or control stimulation was complete, we examined persistent effects for up to 60 minutes.

**Figure 3.**
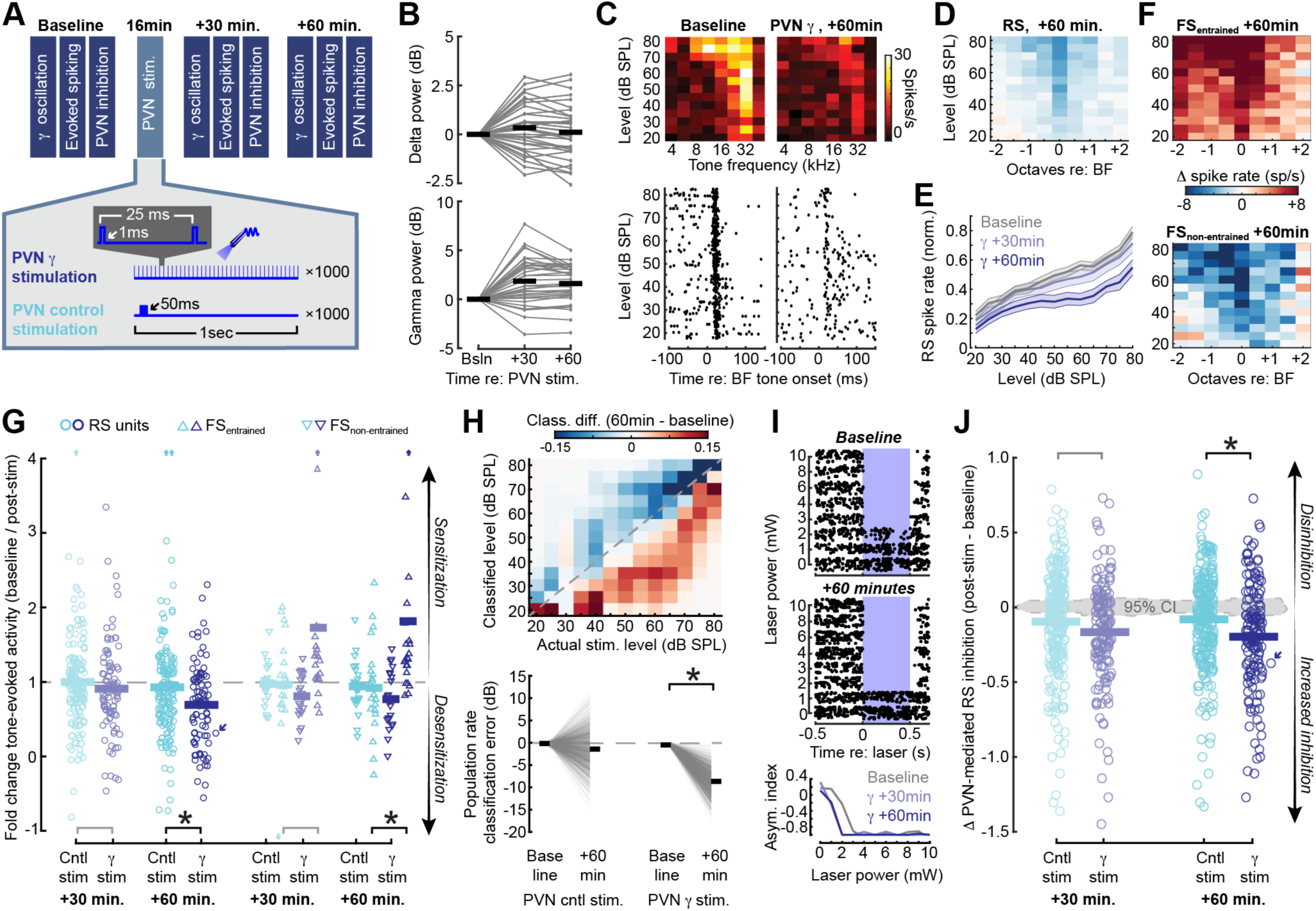
Potentiated PVN-mediated inhibition, gamma rhythms, and sustained reductions in sound-evoked activity after direct activation of PVNs at 40Hz. A) Schematic for PVN γ and PVN control stimulation and recording protocols. B) Steady-state (non-evoked) LFP power in the delta (2-4Hz) and gamma (30-80Hz) range was measured before and after PVN stimulation and expressed as a difference (post-pre). The change in spontaneous gamma power was significantly increased following a single bout of PVN γ stimulation (unpaired t-tests against a population mean of zero, t > 4.5 and p < 1 x 10^-5^ for 30 and 60 minutes) whereas delta oscillations were not significantly changed (t < 0.12, p < 0.17 for 30 and 60 minutes). N = 4 mice / 42 recording sites. C) Example frequency response areas (top) and raster plots (bottom) present changes in frequency tuning and sound-evoked spiking at the best frequency (BF) for a single RS unit. D) Mean reduced sound-evoked firing rates (+60 minutes – baseline) relative to the baseline BF. E) Mean ±SEM Evoked firing rates as a function of sound level normalized to the maximum response during the baseline period. F) As per *D*, but for ChR2+ FS units that were activated during PVN γ stimulation (FS_entrained_, top) or less activated during the period of PVN γ stimulation (FS_non-entrained_, bottom). G) Fold change in evoked firing rate (+30m or +60m / baseline) was calculated for each single unit from a fixed range of effective baseline tone frequencies (BF ± 0.5 octaves, 50-80 dB SPL). Sound-evoked firing rates were persistently increased in FS_entrained_ units and decreased in RS units after PVN γ stimulation but not control stimulation (3-way mixed design ANOVA, main effect for time [F = 380.45, p = 4 x 10^-57^]; main effect for cell type [F = 8.89, p = 2 x 10^-4^]; main effect for stimulation type [F = 2.54, p = 0.11]; cell type x stimulation type interaction [F = 9.77, p = 8 x 10^-5^]; time x cell type x stimulation type [F = 10.17, p = 6 x 10^-5^]; N = 4 PVN γ stimulation mice (n = 91/21/22 units that were significantly sound-evoked at baseline, RS/FS_entrained_ (upward triangles)/FS_non-entrained_ (downward triangles) and 6 PVN control stimulation (n = 163/23/23)). Asterisks indicate significant pairwise differences 60min after PVN γ vs control stimulation for RS units (p = 0.01) and for FS_entrained_ (p = 0.03). Gray lines indicate a non-significant difference (p > 0.07 for both). Upward and downward arrows indicate the occurrence of an outlying value above (upward) or below (downward) the plotted range. Diagonal arrow identifies the unit shown in *C*. *Top:* Confusion matrices depict the change in sound level classification probability between the baseline and 60min after PVN γ stimulation. Classification is based on the population activity rate from 100 RS units. Dashed gray line represents veridical classification. *Bottom*: Decoding was performed on 1000 random samples of 100 RS units from a total pool of n = 311/167 (PVN control/ γ stimulation). Bar = mean. Sound level was significantly biased towards lower, desensitized sound level classification after PVN γ stimulation compared to PVN control stimulation (Mixed design ANOVA, main effect for stimulation type [F = 2205, p < 1 x 10^-60^]; main effect for time [F = 4040.3, p < 1 x 10^-60^]; stimulation type x time interaction [F = 2074.7, p < 1 x 10^-60^]; Asterisk indicates the conjunction of a corrected p value < 0.05 and a large effect size (Hedge’s G) > 2. H) Raster plots show PVN-mediated inhibition from a RS unit measured at baseline and 60 minutes after PVN γ stimulation. PVN-mediated inhibition of spontaneous RS spiking was calculated as an asymmetry index for each eligible RS unit as (Laser_post_ – Laser_pre_ / Laser_post_ + Laser_pre_), where a negative value indicates increased inhibition and a value of zero indicates no difference. Line plots plot the asymmetry indices for the same RS unit shown above at baseline, +30min, and +60min. Diagonal arrow identifies the unit shown in *I*. I) Change in PVN-mediated inhibition after PVN control/ PVN γ stimulation (N = 6/4 mice, 254/147 RS units) measured as the mean asymmetry index after stimulation – mean asymmetry index pre-stimulation, such that negative values represent enhanced inhibition. Gray area represents the 95% confidence interval of PVN-mediated inhibition measured at 0mW, which provides an estimate of measurement noise. PVN-mediated inhibition of RS units is significantly greater 60 minutes after PVN γ stimulation compared to PVN control stimulation (unpaired t-test, t = 3.27, p = 0.001) but is only marginally increased at 30 minutes (t = 1.92, p = 0.06).

For the predicted increase in LFP resting gamma power, the outcome was not entirely clear. We found that LFP gamma power after PVN γ stimulation was significantly increased for electrode contacts within layer 1-6 of A1, while low-frequency delta power was not changed (**Figure 3B**), nor were significant changes observed in either frequency band for contacts in the subcortical white matter (**Figure S1A-B**). However, the same pattern of changes were observed with PVN control stimulation, suggesting that sustained changes in LFP gamma power are not specific to PVN stimulation protocols that elicit strong endogenous gamma rhythms (**Figure S1C**).

We observed a striking and specific effect of PVN γ stimulation on sound-evoked activity, where receptive fields and tone-evoked spiking around the best frequency (BF) in RS units were strongly suppressed 60 minutes after PVN γ stimulation (**Figure 3C-D**). Dampened sound responses appeared more prominent 60 minutes after PVN γ stimulation than 30 minutes and were more pronounced at high sound levels than low sound levels (**Figure 3E**). Suppression of sound-evoked responses in RS units was paralleled by a prominent increase in sound-evoked spiking in PVN FS units that were recruited by the 40Hz laser stimulation (FS_entrained_) while the suppressed sound responses in FS units less activated by PVN γ stimulation resembled RS units (FS_non-entrained_), **Figure 3F**). Whether dampened or enhanced, spike rates were specifically changed near each unit’s BF.^8^

To quantify these effects, we calculated the fold change in sound-evoked spiking for stimuli at or near each unit’s BF in a range of 50-80 dB SPL. Among RS units, we observed that sound-evoked spiking suppression was equivalently weak 30 minutes after stimulation with either protocol but was twice as pronounced with PVN γ stimulation compared with PVN control stimulation by 60 minutes (**Figure 3G, left**). Sound-evoked spiking was increased in PVN neurons, but only in FS_entrained_ units 60 minutes after the cessation of stimulation, and only after PVN γ stimulation, not control stimulation (**Figure 3G, right**). Importantly, the pattern of altered sound-evoked responses here could not be attributed to changes in spontaneous spike rates, which tended to be slightly increased in a manner that was consistent across unit types, post-stimulation time, and stimulation protocols (**Supplemental Figure 1D**). Above, we noted that acute activation of PVNs shifted sound level decoding 17 dB lower than their actual sound level (Figure 1H-J). We also observed a significant ∼10dB shift in population sound level classification 60 minutes after PVN γ stimulation was discontinued but no significant change after PVN control stimulation (**Figure 3H**). Thus, at the level of RS unit population responses in A1, sounds were misclassified as lower than their physical level either when sounds were presented during PVN activation (Figure 1J, left) or for as long as one hour after PVN γ stimulation was discontinued (Figure 3H, right).

Sound-evoked spiking in FS_entrained_ units were approximately doubled 60min after PVN γ stimulation, suggesting that dampened RS responses could arise from enhanced spiking of PVNs. Postsynaptic changes in the RS units could also contribute to dampened sound responses, such as insertion of membrane-bound GABA receptors that could mediate enhanced postsynaptic inhibitory currents.^48^ While we could not directly measure changes in synaptic properties with extracellular recordings, we can gain some insight into inhibitory changes by quantifying changes in the strength of spontaneous RS inhibition when PVNs were activated with laser pulses of increasing power.^49^ In RS units that showed incomplete suppression of spiking following PVN activation at baseline, we observed significantly enhanced PVN-mediated inhibition after PVN γ stimulation (**Figure 3I**).

Inhibitory potentiation was significantly greater with PVN γ stimulation than PVN control stimulation and was observed 60 minutes post-stimulation, but not 30 minutes (**Figure 3J**).

### Behavioral assessment of loudness perception

Our findings presented thus far indicate that A1 PVNs function as a “volume knob” on the population activity rate, shifting the decoded sound level higher or lower than the actual level. Further, our findings identify some “stickiness” in the volume knob, such that we could significantly dampen RS sound responses and shift population-based decoding towards hyposensitivity for at least 60 minutes after a bout of PVN activation that produced endogenous gamma rhythms. A key unsolved question is whether any of these neural changes translate to meaningful changes in loudness perception.

Loudness describes a perceptual characteristic of sound intensity that can be arranged on a scale from quiet to loud.^50^ Because loudness is a subjective psychological state, it can only be directly measured through behavioral reporting of sound perception. We developed a two-alternative forced choice (2AFC) sound level categorization task that conditionally reinforced mice for licking the “soft” spout following presentation of 40-45 dB SPL tones or the “loud” spout for 75-80 dB SPL tones (**Figure 4A, top**). These conditionally reinforced ranges of soft and loud sound levels were presented on 50% of trials and were used to ensure that mice understood the procedural demands of the task and to safeguard against spout choice bias. In the remaining 50% of trials, mice were unconditionally rewarded for categorizing tones of intermediate intensity (50-70 dB SPL) as either soft or loud, as there was no objectively correct answer. This framework ensured that choice was strictly dependent on sound intensity without influencing how mice reported the transition between soft and loud sounds across an intermediate range of levels.

**Figure 4.**
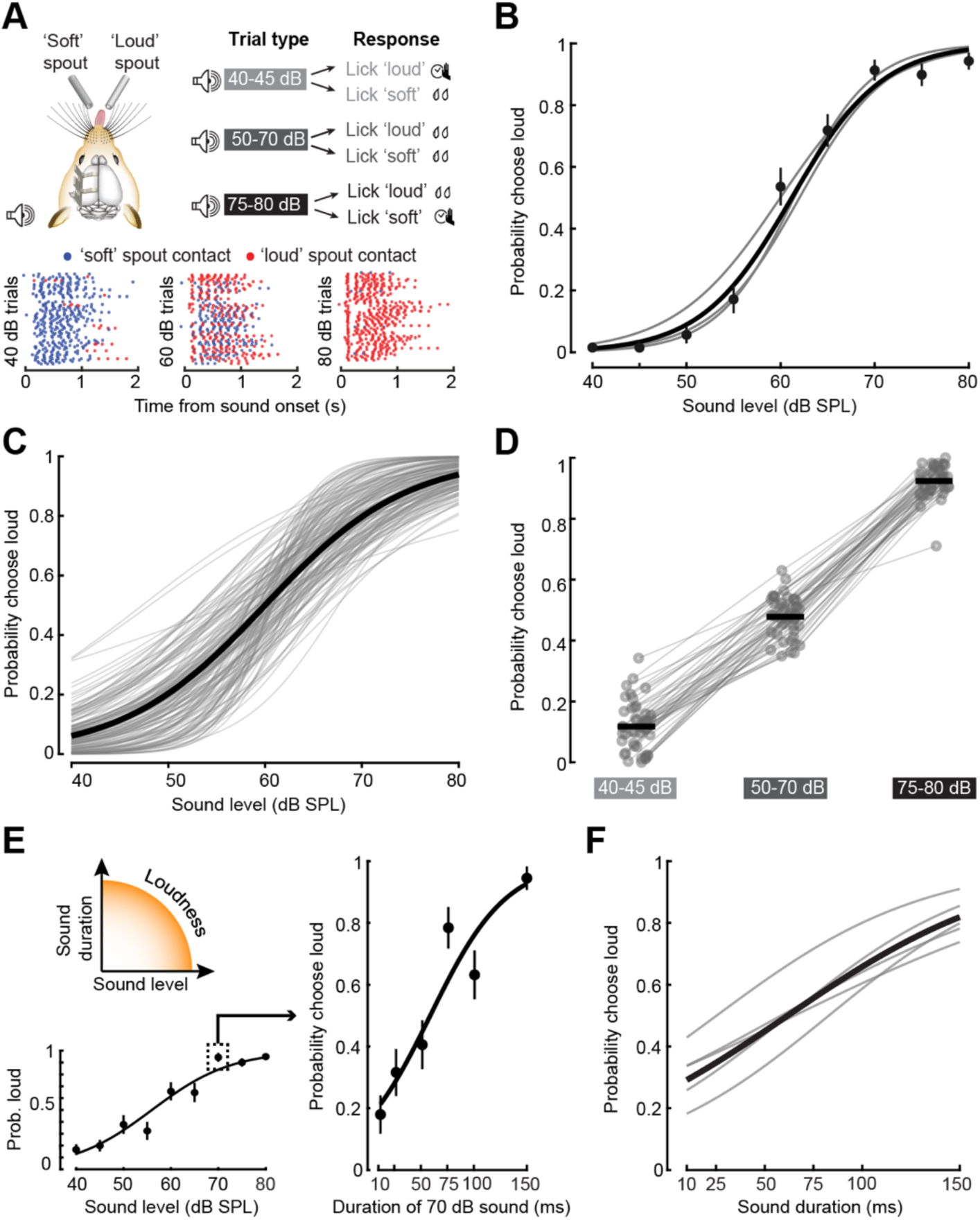
A two-alternative forced choice task to measure loudness perception. A) *Top*, Mice were trained to categorize 11.3 kHz tones as either ‘soft’ or ‘loud’. On approximately 50% of trials, mice were conditionally reinforced for accurate categorization of soft (40-45 dB SPL) and loud (75-80 dB SPL) tones. Mice received water reward regardless of their choice for moderate levels (50-70 dB SPL). *Bottom*, Lick rasters from one example behavioral session for trials at three different intensities. Color represents spout choice. Each row represents a single trial. B) Psychometric functions were fit to raw left/right choice data using binary logistic regression. Fits were applied to the concatenated data (thick black line) drawn from three behavioral sessions (thin gray lines). Error bars = SEM. C) All behavioral choice functions (n = 165 sessions from 47 mice). Thick line represents the grand average of individual sessions. D) Choice probability for all mice in the conditionally reinforced levels (40-45 and 75-80 dB SPL) and moderate unconditionally reinforced probe levels (50-70 dB SPL). Thick lines = sample mean. Each mouse is represented as an individual thin line. E) Schematic illustrates loudness increasing as a function of sound level or sound duration. *Bottom:* Loudness classification for a representative mouse with 150ms tones. *Right:* Loudness classification in the same mouse for a 70 dB SPL tone across a range of shorter durations. Error bars represents the bootstrapped SEM of choice probability at each intensity. F) The probability of reporting a fixed sound level as loud significantly decreased with decreasing sound duration (one-way repeated measures ANOVA, N = 5 mice, F = 46.41, p = 3 x 10^-10^). Solid black line and thin gray lines represent mean and individual mice, respectively.

Mice accurately reported sounds at the low and high ends of the continuum with short-latency licks on the soft and loud spout, respectively (**Figure 4A, bottom**). Mice alternately reported intermediate sound intensities as soft or loud from one trial to the next, yielding a smoothly varying transition in choice probability when averaged across trials (**Figure 4B**). Loudness reporting across behavioral sessions was highly reliable across all mice tested (N = 47; **Figure 4C**), where the probability of categorizing the sound as loud grew monotonically across the range of tested sound levels (**Figure 4D**).

In humans, loudness is the product of sound level and duration, such that a high-intensity sound will be perceived as only moderately loud if it is presented for a short period of time.^51^ To determine whether sound level categorization task captured the perception of loudness, we identified a level that mice reliably reported as loud with the standard 150ms duration and then presented that level across a range of shorter durations (**Figure 4E**). As predicted, the probability that a fixed level tone would be reported as loud was significantly reduced as the tone duration decreased to 10ms, demonstrating that mice were reporting the perception of loudness, not the physical sound level (**Figure 4F**).

### Auditory cortex PVNs regulate loudness perception

Studies have employed physical, pharmacological, or optogenetic inactivation approaches to establish whether the ACtx is a necessary component of a distributed brain network underlying sound perception. The weight of evidence across many studies supports the necessary involvement of the auditory cortex in the discrimination of relatively complex sounds,^52^ sounds presented in background noise,^53^ or sounds requiring temporal integration,^54^ but that ACtx is not necessary for the detection of simple sounds in quiet background.^52^ The electrophysiology experiments described above make a different kind of prediction regarding cortical sound processing and the perception of loudness; not that the ACtx is necessary for the detection of an isolated sound, but rather that ACtx population activity functions as a volume knob for the perceived loudness of that sound.

Specifically, the population decoding experiments suggested that sounds would be perceived as louder than its physical level when population activity was elevated by PVN inactivation (see Figure 1J) but softer than its physical level during PVN activation or following a period of PVN γ stimulation (see Figure 3H).

To test these hypotheses, mice performed the 2-AFC loudness categorization task with the same laser stimulation protocol used in electrophysiology experiments delivered bilaterally to the ACtx on 33% of trials (**Figure 5A**; post-mortem reconstructions of transduced PVN regions are provided in **Figure S2**; targeted expression of ChR2 in PVNs of a subset of mice with the S5E2 enhancer virus is shown in **Figure S3**). Tone detection (i.e., licking either spout) occurred on nearly 100% of trials and was not affected by optogenetic silencing of ACtx activity via PVN activation, consistent with reports that cortical inactivation does not impact tone detection (**Figure 5B**).^52^ However, the probability of reporting moderate and high sound levels as loud was significantly reduced in interleaved PVN activation trials (**Figure 5C-D**). Further, we were able to titrate the reported loudness of a fixed level tone by varying the degree of PVN activation using a range of laser powers (**Figure 5E, top**). We found that a loud sound was increasingly more likely to be reported as soft as the degree of PVN activation was incrementally increased (**Figure 5E, bottom**). As a control for non-specific reporting bias due to laser stimulation, we found that laser activation in mice that expressed GFP rather than ChR2 had no discernable effect on loudness perception (**Figure 5F**). Importantly, increasing ACtx population activity via PVN inactivation had the opposite effect; low and moderate tone levels consistently reported as soft in laser off trials were significantly more likely to be reported as loud during PVN inactivation (**Figure 5G**).

**Figure 5.**
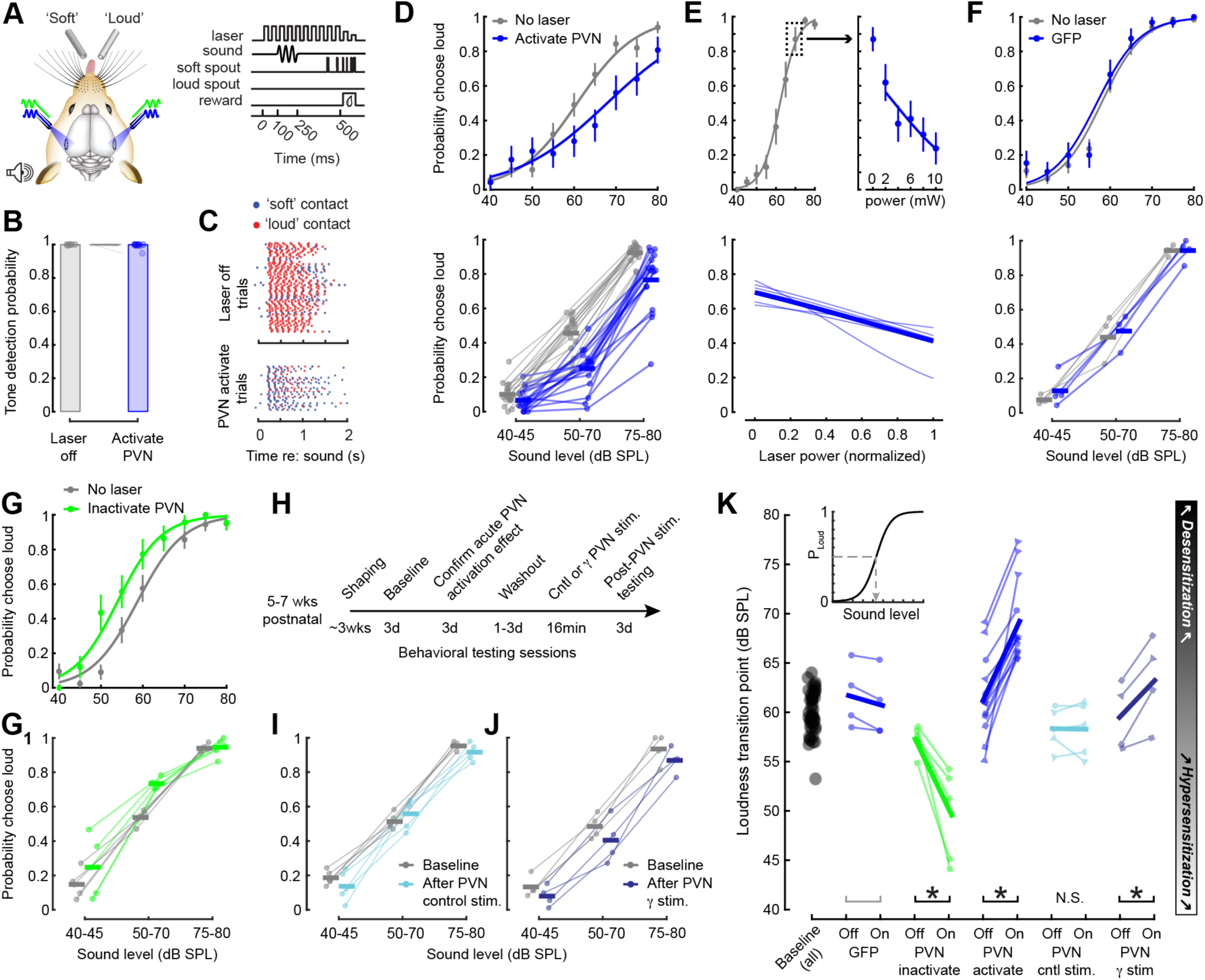
ACtx PVNs regulate loudness perception. A) Cartoon and schematic depict the organization of optogenetic stimulation, sound presentation, lick contact registration, and reward delivery for behavioral studies that combine loudness categorization with bilateral manipulation of PVN activity. B) Cortical silencing via PVN activation did not significantly change the probability of tone detection (N = 14 mice, paired t-test, t = 1.11, p = 0.29). Light gray circles and lines denote individual mice. Bars indicate sample means. C) Lick rasters for a fixed sound level demonstrate a switch in loudness categorization from loud to soft when tones were presented during PVN activation. Each row represents a single trial. D) *Top:* Sound level classification during interleaved laser on and off trials in an example mouse that expressed ChR2 in PVNs. Error bars = SEM. *Bottom:* Choice probability for all mice (N = 14, thin lines) and means (thick horizontal lines) for interleaved laser on and off trials. PVN activation significantly reduced the probability of reporting sound levels as loud (2-way repeated measures ANOVA; main effect for PVN activation [F = 29.9, p = 6 x 10^-5^], level x PVN activation interaction [F = 28.88, p = 7 x 10^-5^]. E) At a fixed sound level reliably perceived as loud, titrating the degree of PVN activation progressively reduced the probability of loud classification in an example mouse (top) and across all mice (thin lines = individual mice; thick line = mean of 5 mice; 1-way repeated measures ANOVA, [F = 9.63, p = 9 x 10^-5^]. F) *Top:* Sound level classification during interleaved laser on and off trials in an example mouse that expressed the control fluorophore GFP in ACtx neurons. Error bars = SEM. *Bottom:* Choice probability for all mice (N = 4, thin lines) and means (thick horizontal lines) for interleaved laser on and off trials. Exciting GFP with blue light had no impact on loudness reporting (2-way repeated measures ANOVA; main effect for PVN activation [F = 1.85, p = 0.27], level x PVN activation interaction [F = 1.35, p = 0.33]. G) *Top:* Sound level classification during interleaved laser on and off trials in an example mouse that expressed Arch in PVNs. Error bars = SEM. *Bottom:* Choice probability for all mice (N = 5, thin lines) and means (thick horizontal lines) for interleaved laser on and off trials. PVN inactivation significantly increased the probability of reporting sound levels as loud (2-way repeated measures ANOVA; main effect for PVN activation [F = 17.4, p = 0.01], level x PVN activation interaction [F = 25.69, p = 0.007]. H) Timeline for experiments that studied persistent changes in loudness classification over several days following 1000 trials of PVN activation concentrated into a 16-minute period. As a positive control for each mouse, we first confirmed that PVN activation during sound presentation reduced the probability of reporting sounds as loud (acute PVN testing) before investigating whether sound level reporting could be stably shifted following control or γ stimulation paradigms. I) Choice probability for each mouse (N = 5, thin lines) and means (thick horizontal lines) from the baseline testing period (gray lines) and post PVN control stimulation (blue lines). A single bout of PVN control stimulation had no impact on loudness reporting (2-way repeated measures ANOVA; main effect for PVN activation [F = 1.19, p = 0.34], level x PVN activation interaction [F = 1.41, p = 0.3]. J) As per *I*, but for PVN γ stimulation. A single bout of PVN γ stimulation significantly reduced the probability of reporting sounds as loud for several days after stimulation (2-way repeated measures ANOVA; main effect for PVN activation [F = 13.57, p = 0.03], level x PVN activation interaction [F = 11.86, p = 0.04]. K) *Inset:* The loudness transition point was defined as the sound level associated with a 0.5 probability of reporting sounds as loud. Individual mice are represented as thin lines and the sample mean as a thick line. Circles denote wildtype mice in which GFP was expressed in ACtx neurons with AAV2/5-hSyn-EGFP or ChR2 was selectively expressed in PVNs with AAV2/5-S5E2-mCherry. Triangles denote transgenic PV-Cre x Ai32 mice that express ChR2 in PVNs. Squares denote PV-Cre mice that express Arch or ChR2 in PVNs with AAV-DIO-ChR2 or AAV-FLEX-Arch. Paired t-tests: GFP (t = 3.02, p = 0.06); PVN inactivation via Arch (t = 4.22, p = 0.01); PVN activation via ChR2 (t = -7.17, p = 2 x 10^-5^); following PVN control activation (t = 0.11; p = 0.92), following PVN γ stimulation (t = -4.0, p = 0.03). Asterisk denotes p value < 0.05. Gray lines = not significant.

These experiments demonstrated that loudness perception can be briefly turned up or down during optogenetic PVN inactivation or activation, respectively. To determine whether mice could be desensitized to loud sounds for extended periods of time, we tracked loudness reporting over a 2-day period following a single 16min bout of PVN γ or control activation (**Figure 5H**). We found that a single bout of PVN control activation had no lasting effect on loudness reporting (**Figure 5I**) while a single bout of PVN γ stimulation produced a shift towards reduced loudness throughout the 2-day post-stimulation test period (**Figure 5J**).

We summarized these effects by calculating the loudness transition point, the sound level where mice exhibited an equal probability of reporting the sound as soft or loud (**Figure 5K, inset**). The loudness transition point was remarkably consistent across the full cohort of unmanipulated control mice (59.97 ± 0.32 dB). Mice expressing GFP in PVNs exhibited no significant change in loudness transition point in laser on vs laser off trials (-1.14 ± 0.37 dB). Mice expressing Arch in PVNs were significantly hypersensitized to loudness during PVN inactivation (-8.15 ± 1.93 dB), while mice expressing ChR2 in PVNs were significantly desensitized to loudness during PVN activation (8.06 ± 1.04 dB). A single intense bout of PVN activation had no effect on loudness perception compared to baseline in the control configuration (0.07 ± 0.75 dB) but induced significant desensitization to loudness when presented at 40Hz (4.08 ± 1.0 dB).

### Noise-induced cochlear damage causes A1 hyper-responsivity and loudness hypersensitivity

These findings show that ACtx PVNs bi-directionally control population coding of sound level and the perception of loudness; inactivating PVNs shifts population decoding and the perception of loudness towards hypersensitivity while activating PVNs shift neural classification and perception towards reduced sensitivity. Further, a single bout of PVN γ stimulation – but not PVN control stimulation – recapitulated the effects of acute PVN activation, in that A1 neuron and behavioral reporting of loudness could be desensitized for an extended period of time after stimulation ends. As a next step, we asked whether these manipulations could be applied to a mouse model of loudness hypersensitivity to restore normal sound level processing.

PVN hypofunction and sensory hypersensitivity have been associated with a wide range of neurological conditions, but we selected to focus on noise-induced cochlear deafferentation because it is a common cause of loudness hypersensitivity that we could initiate in mature animals at a precise timepoint.^55,56^ We caused a controlled sensorineural hearing loss (SNHL) in the high-frequency base of the cochlea by exposing mice to 16-32kHz octave-band noise at 103 dB SPL for two hours (**Figure 6A**). A sham control cohort of mice underwent the same protocol but were exposed to a sound level that did not cause any lasting damage to the cochlea. SNHL noise exposure elevated auditory brainstem response (ABR) thresholds for high frequency tones aligned to the region of sensorineural damage, as confirmed by a significant loss of primary afferent synaptic contacts between inner hair cells and Type-I spiral ganglion neurons in the high-frequency base of the cochlea (**Figure 6C**).

**Figure 6.**
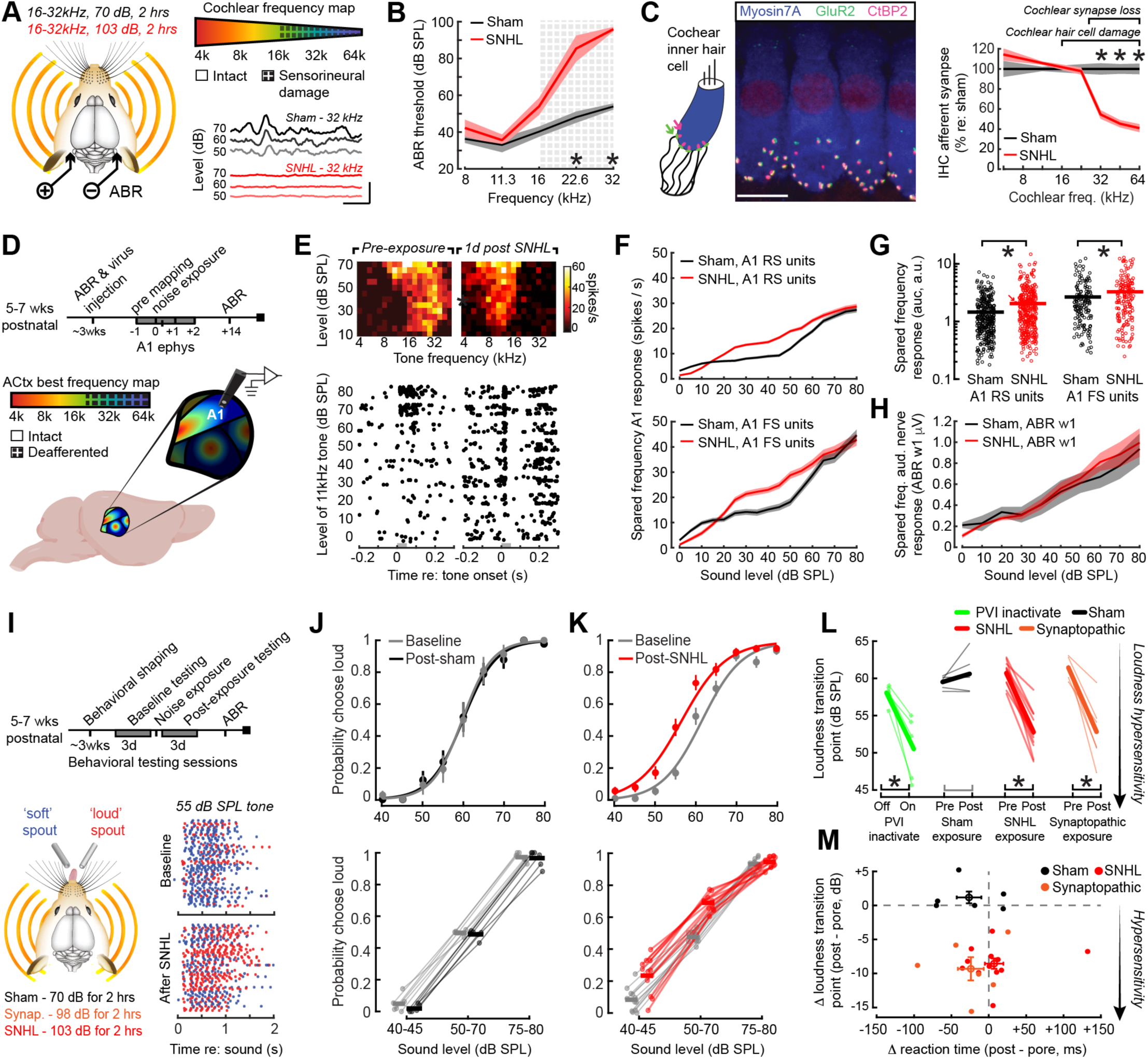
Cortical hyperresponsivity and loudness hypersensitivity to spared sound frequencies following noise-induced sensorineural hearing loss. A) Cartoon illustrates SNHL and sham noise exposure protocols, region of outer hair cell damage and inner hair cell (IHC) synaptic loss in the organ of Corti, and ABR waveforms from an example SNHL and sham mouse elicited by 32kHz tones. Scale bars = 1ms and 1μV. B) Mean ±SEM ABR thresholds were significantly elevated several weeks after SNHL compared to sham exposure, particularly at high frequencies (one-way ANOVA, N = 5/6 sham/SNHL; main effect for group [F = 25.88, p = 0.0007]) Asterisks indicate significant differences with post-hoc pairwise contrasts (p < 0.006). C) *Left*, cochlea immunostained for anti-CtBP2 and anti-GluR2a reveal presynaptic ribbon and post-synaptic glutamate receptor patches on inner hair cells (IHC’s). *Right*, SNHL exposure caused a permanent reduction of IHC synapses in the high frequency region compared to sham-exposed cochleae (Mixed model ANOVA with Group as a factor and Frequency as a repeated measure: Frequency x Group interaction, F = 22.0, p = 5 x 10^-11^). Asterisks denote significant differences between sham and noise exposure with post hoc pairwise comparisons (p < 0.005). Scale bar is 10 µm. D) Experimental timeline for A1 electrophysiology recordings in noise-exposed mice. Lateral view of the mouse brain depicts tonotopic gradients of various fields within the mouse ACtx, highlighting the deafferented high-frequency region of A1, where recordings were made in SNHL mice. E) *Top:* Frequency response areas from two RS units recorded in the high-frequency region of the A1 tonotopic map before and after SNHL. *Bottom:* Rasters from the same units display sound-evoked spiking elicited by 11.3kHz tones of varying levels. Gray bars denote tone duration. F) Mean ±SEM spike rate-level functions in SNHL RS (top, N/n = 6/376) and FS (bottom, n = 172) units are both significantly greater than sham RS (N/n = 5/339) and FS (n = 155) units (Two-way mixed design ANOVA, level x group interaction, F > 4.75, p < 2 x 10^-9^ for both). G) Sound-evoked A1 responses to 11.3kHz spared frequency tones from the same units in *F* are significantly increased in SNHL mice compared to sham (Mixed design ANOVA, main effect for exposure group [F = 12.17, p = 4 x 10^-4^]). Asterisks denote significant pairwise contrasts for RS and FS, respectively (p = 4 x 10^-5^ and 4 x 10^-4^, respectively). Each single unit is depicted as a circle. Horizontal bar = mean. Diagonal arrow illustrates the RS unit shown in *E*. H) Sound-evoked ABR wave 1 responses to 11.3kHz spared frequency tones were not different between SNHL/Sham (N = 6/5) (Mixed design ANOVA, main effect for exposure group [F = 0.05, p = 0.83]). I) Experimental timeline for the 2AFC behavioral studies that tracked sound level categorization before and after exposure to sham, synaptopathic, and SNHL noise levels. Lick rasters display single trial categorization for behavioral sessions from the same mouse before and after SNHL at a moderate, unconditionally reinforced sound level. J) *Top:* Sound level classification before and after sham exposure in an example mouse. Error bars = SEM. *Bottom:* Choice probability for all sham mice (N = 6, thin lines) and means (thick horizontal lines) from the baseline and post-exposure period. Sham exposure had no impact on loudness reporting (2-way repeated measures ANOVA; main effect for Time [F = 2.85, p = 0.15], Level x Time interaction [F = 1.65, p = 0.26]. K) As per *J*, *Top:* an example SNHL mouse. *Bottom:* Choice probability for all SNHL mice (N = 12, thin lines). SNHL exposure significantly increased the probability of reporting sounds as loud, particularly at low and moderate sound intensities (2-way repeated measures ANOVA; main effect for Time [F = 107.3, p = 6 x 10^-7^], Level x Time interaction [F = 99.15, p = 8 x 10^-7^]. Pairwise contrasts identified significant differences at 40-45 and 50-70 level ranges (p < 2 x 10^-5^ for both). L) Loudness transition points for individual mice (thin lines) and sample means (thick line). PVI inactivation data with Arch are replotted to facilitate direct comparison to noise exposure. Paired t-tests: Sham (t = 3.02, p = 0.06); SNHL (t = 11.33, p = 3 x 10^-7^); synaptopathic noise exposure (N = 6; t = 5.47, p = 0.003). M) Scatterplot depicts the change in loudness transition point against the change in reaction time for individual mice (circles) and means (open circles, bi-directional error bars = SEM). Reaction time was not significantly changed after noise exposure (One-way ANOVA [F = 1.77, p = 0.2) and no significant correlation was observed between loudness hypersensitivity and reaction time (Pearson r = -0.14, p = 0.52).

To document how a restricted high-frequency cochlear lesion affected cortical sound processing, we first performed electrophysiological mapping to identify the high-frequency region of the A1 tonotopic map one day before SNHL and then continued to perform extracellular single unit recordings from the A1 high-frequency region up to several days after hearing loss (**Figure 6D**). Prior to noise exposure, most RS units were tuned to sound frequencies in the 16-45kHz range with weak, high-threshold responses to mid-frequency tones (**Figure 6E, left**). Days after SNHL, single RS units were tuned to spared mid-frequency tones (**Figure 6E, right**), consistent with a competitive adult cortical map plasticity that has been observed following mechanical, chemical, noise-induced, or natural degeneration of the high-frequency cochlear base.^57^ Looking across all sound-responsive single units, we noted significantly greater growth of cortical RS and FS responses to spared 11.3 kHz tones in SNHL compared to sham mice (**Figure 6F-G**). Peripheral neural responses to 11.3kHz tone bursts, as measured by the amplitude of ABR wave 1, were not different between sham and SNHL mice (**Figure 6H**) confirming prior reports that the over-representation of spared sound frequencies following restricted cochlear lesions reflects a compensatory plasticity at higher stages of the central auditory pathway.^58,59^

Next, we investigated whether cortical hyper-responsivity to spared mid-frequency tones after SNHL was associated with behavioral loudness hypersensitivity. Because SNHL could be induced at a precise day, we were able to use each mouse as their own control and study the development of sound sensitivity to mid-frequency tones near the edge of the cochlear lesion before and up to two weeks after noise exposure (**Figure 6I**). Sham-exposed mice did not show a significant change in the probability of reporting 11.3 kHz tones as loud after noise exposure (**Figure 6J**), whereas SNHL mice exhibited loudness hypersensitivity, as measured both by the a significantly greater probability of reporting low and moderate sound intensities as loud (**Figure 6K**) and a -8.55 dB reduction in their loudness transition point, almost exactly matching the loudness hypersensitivity produced by optogenetic PVN inactivation (**Figure 6L**).

Noise intensities high enough to induce a permanent ABR threshold shift damage cochlear outer hair cells in addition to the loss of cochlear afferent neural synapses onto inner hair cells described above. Outer hair cell damage can reduce the specificity of cochlear frequency tuning, implying that hypersensitivity to spared mid-frequency tones might be simply attributable to a change in peripheral sound transduction.^60^ We ruled out a contribution of peripheral biomechanical changes by also including a cohort of synaptopathic mice exposed to a more moderate sound intensity that damaged cochlear afferent synapses with minimal effects on outer hair cells or ABR thresholds (**Supplemental Figure 4**).^61^ We observed significantly increased loudness sensitivity in mice with purely synaptopathic lesions that was on par with the more extreme SNHL exposure group, implying that central compensatory plasticity following cochlear denervation was the root cause of loudness hypersensitivity after noise-induced hearing loss (**Figure 6L**). Previous studies have modeled loudness sensitivity with systemic administration of highly concentrated salicylate, the active ingredient in aspirin.^62^ Rather than measure loudness perception directly, prior work has relied on a behavioral proxy for loudness sensitivity via a reduced reaction time in a Go-NoGo task.^62^ Here, we found that mice with explicitly measured loudness sensitivity following SNHL or synaptopathic noise exposure did not exhibit reduced reaction times, nor did we note any significant correlation between the shift in loudness transition point and change in reaction time, demonstrating that reaction time is not a stand-in for loudness reporting with models of noise-induced sound sensitivity (**Figure 6M**).^63^

### ACtx PVN function is both a cause and solution for disordered loudness sensitivity after SNHL

Prior work from our lab and others have implicated disinhibition particularly from PVNs as an underlying cause of ACtx hyperactivity after sudden hearing loss.^23,49,64^ Here, we reasoned that if PVN activation enhanced endogenous gamma power (Figure 2F), dampened cortical responses to sounds of increasing level (Figures 1H and 3E), reduced loudness sensitivity (Figure 5K), and increased PVN-mediated inhibition (Figure 3I), then SNHL might be associated with the opposite pattern of effects; namely, that hyper-responsivity (Figure 6F) and loudness hypersensitivity (Figure 6L) might be associated with reduced gamma oscillations and reduced PVN-mediated inhibition. We tested these predictions in the same extracellular recording sessions described above (**Figure 7A**). Filtering the columnar recordings for LFPs rather than spikes confirmed a significant reduction in gamma power in SNHL mice compared with sham but no change in the subcortical white matter gamma nor in delta frequencies (**Figure 7B**). In normal hearing mice, spontaneous RS activity rates were suppressed by PVN activation at laser powers as low as 1mW and was completely quenched above 5mW (**Figure 7C-D**). After SNHL, RS unit spiking was significantly less sensitive to optogenetic PVN activation (**Figure 7E**). However, FS units directly activated by optogenetic activation exhibited comparable recruitment functions in sham and SNHL mice (**Figure 7F-H**). These findings suggest that disinhibition of A1 RS units was not attributable to a reduced efficiency of direct PVN activation but instead arose from pre-or postsynaptic modifications to synaptic inhibition in local A1 circuits.^65^

**Figure 7.**
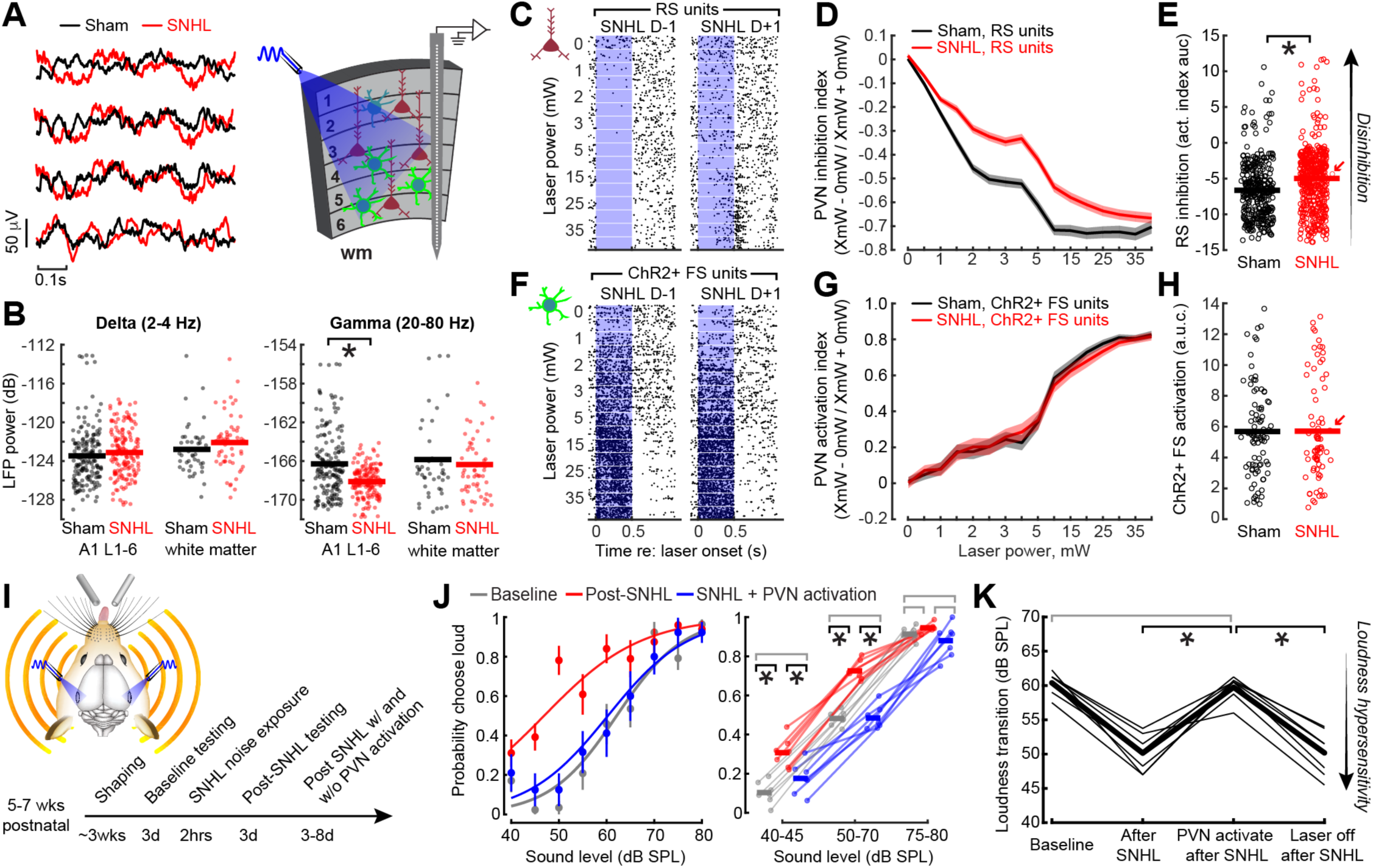
ACtx PVN activation is sufficient to rescue loudness hypersensitivity after SNHL. A) Cartoon illustrates LFP waveforms from electrodes in the upper, middle, and deep layers of A1 as well as subcortical white matter. Recordings of LFP and single unit spiking were performed with and without optogenetic activation of PVNs transduced with ChR2. B) Steady-state (non-evoked) LFP power in the gamma (30-80Hz) and delta (2-4Hz) range was measured after SNHL or sham exposure. The change in spontaneous gamma power in A1 was significantly reduced in SNHL mice (unpaired t-test [t = 6.27, p = 2 x 10^-9)^ but no significant changes were observed in the subcortical white matter or in the delta band (t < 0.76 and p > 0.19 for all comparisons). N = 6/5 mice and 26/19 recording sites for SNHL/sham. C) Rasters present spontaneous spike events for single RS units recorded in the high-frequency region of the A1 tonotopic map the day before and day after SNHL. PVN activation was titrated by varying the laser power across a 0-40 mW range. D) Mean ±SEM PVN-mediated inhibition of RS units was calculated with an asymmetry index where the spike rate within the 0.5s laser period were compared to the 0mW control. Negative values reflect reduced spiking relative to the no-laser 0mW condition. PVN-mediated inhibition of RS units across laser powers was significantly reduced in SNHL mice compared to sham controls (two-way mixed design ANOVA on 471/343 RS units in 6/5 SNHL/sham mice, laser power x exposure group interaction [F = 5.14, p = 3 x 10^-10^]). E) Area under the laser power x asymmetry index curve shown in *D* calculated for each RS unit. Horizontal line = mean. Asterisk indicated that SNHL units were significantly less inhibited by PVN activation (unpaired t-test [t = 5.47, p = 0.02]). Circles denote each RS unit. Diagonal arrow identifies RS unit shown in *C*. F) As per *C*, but for two ChR2+ FS units directly activated by the laser. G) As per *D*, but for FS unit directly activated by the laser in SNHL (N/n = 6/74) and Sham (N/n = 5/88) mice, where positive values identify direct optogenetically elicited spiking. The increase in laser-evoked spike rate in ChR2+ PVNs did not differ between SNHL and Sham mice (two-way mixed design ANOVA, Power x Exposure interaction [F = 0.36, p = 0.98]). H) As per *E,* area under the asymmetry index curve for the FS units in *G* were not different between SNHL and sham units (unpaired t-test [t = 4 x 10^-9^, p = 1.0]). Diagonal arrow identified example unit in *F*. I) Experimental timeline for 2AFC behavioral studies tracking sound level categorization before and after SNHL with and without bilateral PVN activation. J) *Left:* Sound level categorization in an example hypersensitive SNHL mouse was restored to baseline during PVN activation. Error bars = SEM. *Bottom:* Choice probability for SNHL mice (N = 7) were measured at pre-exposure baseline, after SNHL, and in a third phase in SNHL mice with interleaved PVN activation and laser off trials. Thick horizontal lines = mean; individual mice are shown as thin lines. from the baseline and post-exposure period. SNHL exposure significantly increased the probability of reporting sounds as loud, particularly at low and moderate sound intensities (2-way repeated measures ANOVA; main effect for Time [F = 107.3, p = 6 x 10^-7^], Level x Time interaction [F = 99.15, p = 8 x 10^-7^]). Asterisks and black lines indicate significant pairwise differences with Bonferroni-Holm correction for multiple comparisons (p < 0.02 for all); gray lines indicate a non-significant difference (p > 0.06 for all). K) Loudness transition points for individual mice (thin lines) and sample means (thick line) in SNHL mice (N=7). Asterisks and black lines indicate significant pairwise differences (paired t-tests with Bonferroni-Holm correction for multiple comparisons, p < 0.002 for all); gray line indicates a non-significant difference (p = 0.56).

These findings suggested that ACtx PVN-mediated disinhibition could be a root cause of auditory hyperresponsivity and hyperacusis. On the other hand, subcortical stages of sound processing also express hyperactivity after noise-induced hearing loss,^66^ and other inhibitory cell types in the cortex^67^ and elsewhere^68^ are also affected by hearing loss. Because PVNs regulate the perception of loudness, we reasoned that direct activation of PVNs in SNHL mice with sound sensitivity would be sufficient to restore normal loudness perception (**Figure 7I**). As predicted, ACtx PVN activation immediately reversed loudness hypersensitivity and restored baseline loudness perception in SNHL mice (**Figure 7J**). Importantly, this rescue was temporary; it only worked when the laser was on. The loudness transition point reverted to post-noise exposure hypersensitivity on interleaved laser-off trials (**Figure 7K**).

### Sustained reversal of hyperacusis following a single bout of ACtx PVN γ stimulation

Our findings suggest that loudness sensitivity after SNHL arose from compensatory plasticity triggered by the damage of cochlear hair cells and the loss of primary afferent neural synapses. As this peripheral degeneration is a permanent feature, the implication is that brain circuits will revert to a mode of hyperactivity and loudness sensitivity as soon as optogenetic stimulation or drug administration is discontinued. Alternatively, ACtx population activity regulates the perception of loudness and is not strictly beholden to bottom-up inputs, as evidenced by persistent suppression and loudness desensitization following PVN γ stimulation (Figure 4) or persistent hypersensitivity to sounds paired with cortical acetylcholine release.^8^ This raises the possibility that single bout of intense ACtx PVN γ stimulation might be sufficient to desensitize the noise-exposed brain and restore normal loudness perception even though the action is local and inner ear damage persists.

To address this point, we performed an extended course of behavioral loudness testing in SNHL mice (**Figure 8A**). Following SNHL, three days of behavioral testing confirmed that all mice exhibited loudness hypersensitivity to presentation of spared, mid-frequency tones that bordered the cochlear lesion (**Figure 8B**). Mice that subsequently received no treatment or PVN control stimulation exhibited stable hypersensitivity, as measured both by the elevated probability of classifying moderate sound intensities as loud (**Figure 8C, left and middle**) and a reduced transition point between soft and loud sounds (**Figure 8D, left and middle**). However, following PVN γ stimulation, the probability of reporting moderate sound intensities as loud and the loudness transition point both reverted to baseline levels (**Figure 8C-D, right**). Extended testing in SNHL mice demonstrated that loudness sensitivity could be reversed for approximately 1 week after a single 16-minute bout of PVN γ stimulation before reverting to post-exposure levels (**Figure 8E**). These findings highlight the essential role of ACtx PVNs in loudness perception and demonstrate that rekindling their local circuit connectivity can reverse loudness hypersensitivity, even if the initial triggering condition (e.g., cochlear deafferentation) remains uncorrected.

**Figure 8.**
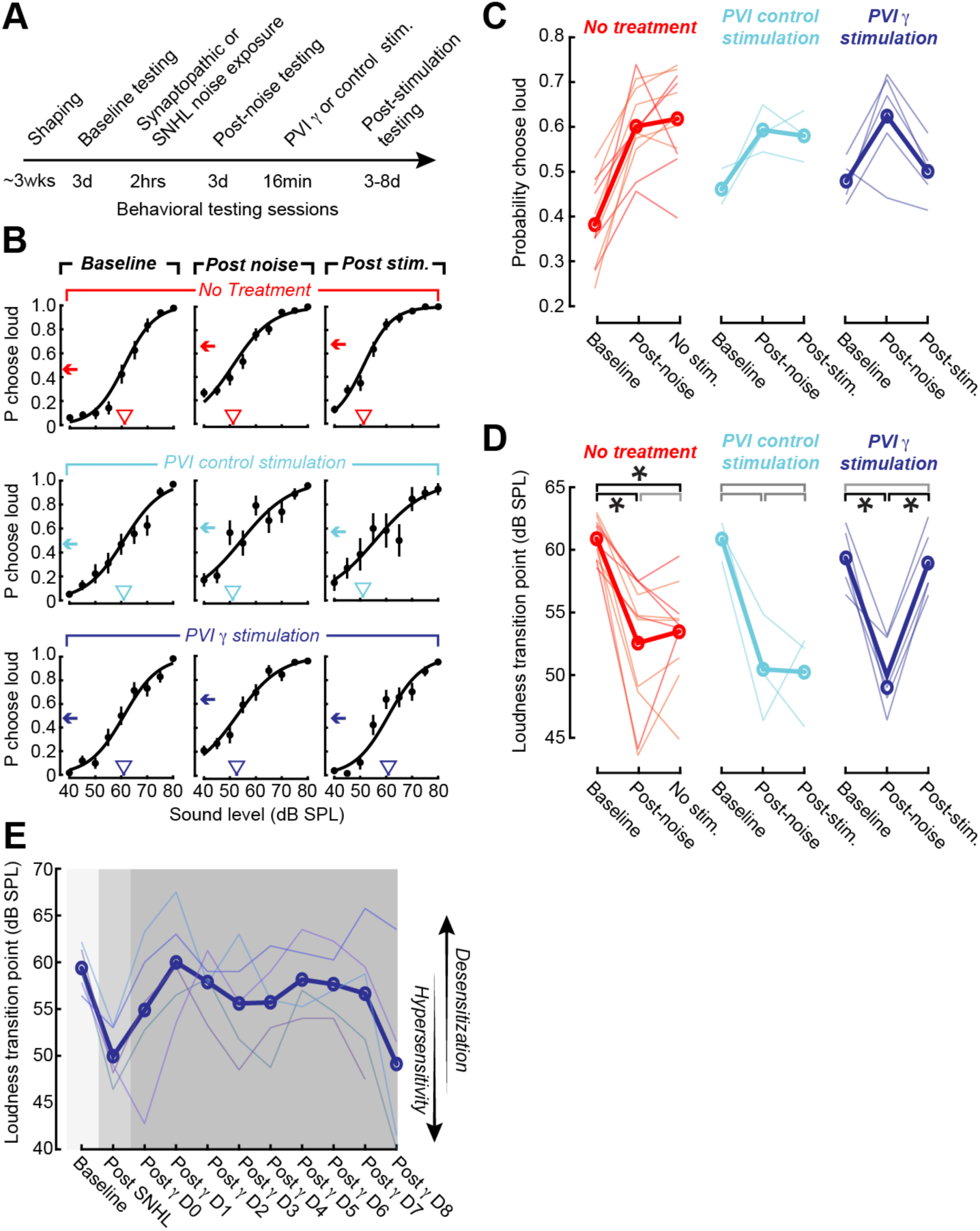
A single bout of PVN γ stimulation reverses loudness hyperacusis for one week. A) Timeline for experiments that seek to induce a sustained reduction in loudness hyperacusis after noise-induced hearing loss with a single bout of PVN activation. B) Sound level categorization in three example mice measured at baseline, after noise exposure, and after a stimulation period. Error bars = SEM. Arrow indicates the mean probability of selecting the loud spout across the range of unconditionally reinforced moderate sound levels (50-70 dB SPL). Downward triangle = loudness transition point. C) Mean choice probability within the unconditionally reinforced range of moderate sound levels for mice that received no stimulation after the post-exposure test session (N = 5/6, SNHL/synaptopathic exposure level, red/orange), PVI control stimulation (N = 3 SNHL), or PVI γ stimulation (N=5 SNHL). The change in choice probability across sessions was significantly dependent on stimulation group (mixed design ANOVA, main effect for session [F = 29.5, p = 6 x 10^-8^]; session x group interaction [F = 5.99, p = 0.001]). Thick line = sample mean. D) As per *C*, but for the loudness transition point. The change in choice probability across sessions was significantly dependent on stimulation group (mixed design ANOVA, main effect for session [F = 39.6, p = 3x 10^-9^]; session x group interaction [F = 5.15, p = 0.003]). Thick line = sample mean. Asterisks and black lines indicate significant pairwise differences (paired t-tests with Bonferroni-Holm correction for multiple comparisons, p < 0.04 for all); gray lines indicate a non-significant difference (p > 0.06 for all). E) Loudness transition point across an extended period of testing in five SNHL mice that received PVN γ stimulation. Thin lines represent individual mice. Thick line = sample mean.

## Discussion

Our experiments addressed how cortical neurons represent the physical quantity of sound level to support the perceptual quality of loudness. We found heterogeneous single neuron encoding of sound level that summed together to form a linear population rate code that could accurately decode sound level without considering the identity or spike times of individual neurons (Figure 1). We observed that 40 Hz PVN activation was especially effective at entraining RS spiking at gamma frequencies (Figure 2) and produced a sustained period of dampened sound responsiveness and enhanced PVN-mediated inhibition (Figure 3). Whether sounds were presented during optogenetic manipulations of PVN activity or up to an hour after PVN γ stimulation, our neurophysiological experiments likened PVN activity to a volume knob, adjusting the sound level decoded from cortical population activity by as much as 25 dB from the physical stimulus level. To relate these observations to loudness perception, we developed a 2AFC loudness classification task (Figure 4) and discovered that not only could the perceived boundary between loud and soft be bi-directionally adjusted by PVN activation or inactivation, but that a single 16-min bout of PVN γ stimulation desensitized normally hearing mice to loud sounds for several days (Figure 5). We then applied these insights in mice with restricted cochlear damage that exhibited behavioral loudness hypersensitivity and neural hyper-responsivity in deafferented regions of A1 (Figure 6). We found that PVN-mediated feedforward cortical inhibition was depressed in mice with noise-induced cochlear damage, but that behavioral loudness hypersensitivity could be temporarily reversed during periods of direct PVN activation (Figure 7). Finally, we observed that normal loudness sensitivity could be rescued in mice with permanent cochlear damage for as long as one week following a single 16-minute bout of PVN γ stimulation, highlighting ACtx PVNs as both a source and solution for perceptual loudness hypersensitivity (Figure 8).

### Possible mechanisms underlying sound dampening, PVN potentiation, and loudness desensitization

ACtx receptive fields express a rapid, specific, and persistent reorganization when sounds of a fixed frequency and level are repeatedly paired with direct stimulation of the cholinergic basal forebrain,^8,69,70^ noradrenergic locus coeruleus,^71^ dopaminergic ventral tegmental area,^72^ or oxytocinergic lateral hypothalamus. ^73^ The cortical over-representation of sounds paired with neuromodulatory stimulation has been associated with improved behavioral detection or discrimination of the corresponding sound stimulus and is supported by an initial reduction in intracortical synaptic inhibition onto excitatory pyramidal neurons that precedes a restructuring and rebalancing of the reorganized receptive field to align with the paired sound frequency.^74^

Here, we sought to expand on this history of targeted stimulation studies in ACtx by identifying a protocol to broadly dampen the cortical response to high-intensity sounds. PVN γ stimulation selectively dampened high-intensity stimuli near the BF without affecting spontaneous activity rates (Figure S1), but took longer to emerge after the stimulation period than the rapid reorganization reported with sound and neuromodulator pairing, a timecourse that may suggest a transcription-dependent process.^75^ Dampened sound responses in RS units were associated with stronger PVN-mediated inhibition of spiking and enhanced sound responses in FS PVN units were entrained during the preceding 40Hz stimulation period. These findings suggested that PVN γ stimulation might enlist a combination of pre-synaptic modification in PVN GABA release and/or post-synaptic changes in pyramidal neuron GABA receptors that would collectively potentiate intracortical inhibition and dampen responses to sound-evoked thalamocortical inputs.^48,76,77^

One limitation of our work is that the behavior and neurophysiology approached used here do not provide direct evidence for a mechanism underlying long-term dampening of cortical sound responses and loudness desensitization. An underlying mechanism could be as straightforward as a potentiation of inhibitory synaptic currents but there is also reason to believe that it could be far more complicated, requiring many additional studies to fully resolve. For instance, modifications of PVN-PVN synapses may also play a role, as the strength and number of these synapses is bidirectionally regulated by PVN activity levels through cell autonomous regulation of production of the neuropeptide VGF.^78^ PVN activation can also increase PVN activity and inhibit PyrN activity millimeters away from the stimulation site.^79^ As such, the effects of PVN γ stimulation may also be mediated by network level effects due to electrical coupling between PVNs^80^ or dense recurrent connectivity between cortical regions.^28^

There are also reasons to speculate that auditory desensitization after PVN γ stimulation may not be directly mediated by synaptic plasticity, as bouts of 40Hz PVN or sensory stimulation have been shown to cause increased cerebral blood flow,^43,81^ neurotrophic factor expression,^82^ neuropeptide release,^81^ glymphatic clearance,^81^ and decreased inflammatory responses.^43^ Importantly, these effects are linked to stimulation-evoked changes in neural activity and not the direct vasodilation of cortical arterioles by light stimulation,^43^ consistent with the fact that no changes were observed in our PVN control stimulation group which received 25% more light exposure over the course of the stimulation compared to the γ stimulation group (50 vs. 40 ms of blue light per second).

Regardless of the precise molecular mechanisms which mediate suppressed ACtx PyrN sound-evoked spiking, our results show that PVN activation protocols impart benefits for sensory coding and perception in the normal and disordered brain. While previous work has primarily linked gamma-band activity to moment-to-moment fluctuations in sensory encoding,^83^ our work suggests that the effects of brief periods of gamma-band entrainment on sensory processing should also be considered on the scale of hours and days. In our study, the choice of 1000s of PVN g stimulation at 10 mW was relatively arbitrary. Future work will be needed to delineate the precise conditions (e.g. duration, intensity, etc.) sufficient to produce suppressed sound evoked activity or to further prolong the effects of γ stimulation.

### Implications for the mechanisms and treatment of sensory disorders

Physical sensory cues are transduced by peripheral sense organs and isomorphically represented by specialized neural circuits distributed throughout the brainstem and midbrain. At the level of the cortex, local inhibitory circuits integrate bottom-up sensory evidence with top-down inputs encoding internal state, sensory predictions, and memory to produce comparatively abstract stimulus representations that underlie perceptual awareness and support adaptive behaviors.^84^ In this regard, cortical inhibitory cell types are key mediators of bottom up and top-down cues that shape the real-time perception of the sensory environment^85^ and regulate long-term activity- and experience-dependent plasticity.^86^

The central role of cortical microcircuits in shaping perceptual representations has important implications for sensory disorders. On the one hand, organisms with normally functioning systems for sensory transduction and encoding can nonetheless have catastrophically disordered perception when defects in local or long-range cortical circuits distort sensory evidence to produce sensory overload^25^ or the hallucination of phantom events.^87^ These circuit defects can arise from inherited susceptibilities and manifest early in development or can be delayed until adulthood, as evidenced by neurodevelopmental disorders or schizophrenia, respectively.^18,20^

By the same logic, peripheral damage or intense peripheral stimulation that would normally produce pain, sensory overload, or sensory hallucinations (e.g., phantom limb, tinnitus) can be perceptually cancelled out through manipulations that reverse aberrant activity at the level of the cortex. This idea is relatively well studied in the context of pain, where optogenetic or chemogenetic manipulation of inhibitory neurons in the somatosensory cortex,^88^ can impart anti-nociceptive effects on the perception of erstwhile painful stimuli by overriding aberrant activity patterns in dense networks of descending corticofugal neurons.^89,90^ In the context of hearing, auditory corticofugal neurons develop marked hyperactivity after noise-induced hearing loss,^91^ underlie learned auditory threat associations^92^ and play a central role in auditory hallucinations.^93^ Monitoring and manipulating auditory corticofugal neurons either directly or indirectly via inhibitory circuit manipulations may provide new insights into the mechanisms and therapeutic considerations for common auditory hyperactivity disorders, such as tinnitus and hyperacusis.

Prior work emphasized that cognitive impairment, perceptual distortions, and pain arising from genetic defects or noxious peripheral inputs can be perceptually renormalized through cortical circuit manipulations, but doing so requires the continuous manipulation of cortical cell types.^25,26,88–90,94^ Here, we confirmed that activating PVNs with a typical optogenetic stimulation protocol rescued loudness hypersensitivity when the laser was on but that mice immediately reverted to loudness hypersensitivity on interleaved laser off trials. PVN γ stimulation provided an intriguing exception to the rule and suggested that perceptual attenuation of intense, painful, or phantom sensory events need not require a continuous, ongoing manipulation of cortical activity. Rather, by exploring particular activation protocols or cell types, it may be possible to impose longer lasting normalization of cortical perceptual representations despite irreversible abnormalities in peripheral sensory input, as exemplified here through 40Hz PVN activation.

## Acknowledgements

These studies were supported by NIH grants DC009836 (DP), DC015857 (SK), DC019128 (EK) and NIH fellowship DC018974 (MM). We thank Y. Watanabe and O. Stevenson for their contributions to animal surgeries and L. Casey for contributions to cortical histology. We thank E. Smith and C. Rutagenwa for assisting with cochlear histology experiments. The Massachusetts Eye and Ear Gene Transfer Vector core produced multiple AAV vectors used in this study.

## Author contributions

Conceptualization, K.K.C., B.A., M.R.M. and D.B.P.; Methodology, K.K.C., M.R.M., K.E.H.; Investigation, K.K.C, B.A., M.R.M., K.S.S., C.K., D.S., D.N., J.Z.; Software, K.K.C., M.R.M.,B.A., K.E.H.; Formal Analysis, K.K.C., B.A., M.R.M.; Data Curation, K.K.C., B.A., M.R.M., J.Z., K.S.S. ; Visualization, K.K.C., B.A., M.R.M., D.B.P., Writing – Original Draft, D.B.P. and K.K.C.; Writing – Review & Editing, K.K.C. and D.B.P; Resources, D.B.P; Supervision, D.B.P.,E.D.K.,S.G.K.; Funding Acquisition, D.B.P., E.D.K., S.G.K., M.R.M.

## Methods

### Key resources table

| REAGENT or RESOURCE | SOURCE | IDENTIFIER |
| --- | --- | --- |
| Bacterial and virus strains |  |  |
| AAV5-Ef1a-DIO hChR2(E123T/T159C)-EYFP | Addgene | 35509 |
| AAV9-Flex-Arch-GFP | Addgene | 22222 |
| AAV2/5-S5E2-ChR2-mCherry | Mass Eye and Ear Gene Transfer Vector Core | Addgene 135634 |
| AAV2/5-Syn-GFP | Mass Eye and Ear Gene Transfer Vector Core | Addgene 58867 |
| Mouse anti-PV Monoclonal IgG1 | Swant | PV235 |
| Alexa Fluor 488 Donkey Anti-Goat IgG | Abcam | ab150129 |
| Rabbit anti-myosin VIIa | Proteus Biosciences | 25-6790 |
| Mouse IgG2a anti-GluA2 | Millipore | MAB397 |
| Mouse IgG1 anti-CtBP2 | BD Biosciences | 612044 |
| AlexaFluor 647 donkey anti-rabbit | ThermoFisher | A-31573 |
| Alexa Fluor 568 goat anti-mouse IgG1 | ThermoFisher | A-21124 |
| Alexa Fluor 488 goat anti-mouse IgG2a | ThermoFisher | A-21131 |
| Deposited data |  |  |
| Source data | This paper; Mendeley | TBD |
| Experimental models: Organisms/strains |  |  |
| Mouse: C57BL/6J | The Jackson Laboratory | RRID:IMSR_JAX:000664 |
| Mouse: B6N(Cg)-Cdh23tm2.1Kjn/Kjn | The Jackson Laboratory | RRID:IMSR_JAX:018399 |
| Mouse: B6.129P2-Pvalbtm1(cre)Arbr/J | The Jackson Laboratory | RRID:IMSR_JAX:017320 |
| Mouse: B6.Cg-Gt(ROSA)26Sortm32(CAG-COP4*H134R/EYFP)Hze/J | The Jackson Laboratory | RRID:IMSR_JAX:024109 |
| Mouse: B6;129S6-Chattm2(cre)Lowl/J | The Jackson Laboratory | RRID:IMSR_JAX:006410 |
| Software and algorithms |  |  |
| Labview 2020 | National Instruments | https://www.ni.com/en-us/shop/labview.html |
| Synapse | Tucker-Davis Technologies | http://www.tdt.com/component/synapse-software/ |
| Kilosort 2.0 | Github | https://github.com/MouseLand/Kilosort |
| AP-Histology | Github | https://github.com/petersaj/AP_histology |
| ABR peak analysis | Github | https://github.com/EPL-Engineering/abr-peak-analysis |
| Matlab 2021b | Mathworks | https://www.mathworks.com/products/matlab.html |
| Original code for data analysis | Github | https://github.com/Plastic-Polley-Lab/TBD |
| Other |  |  |
| PXI controller | National Instruments | PXI-8840 |
| Silicone recording probes | Cambridge Neurotech | H2, H3 |
| Neurodigitizer and preamplifier | Tucker-Davis Technologies | PZ5 |
| Data processor and real-time controller | Tucker-Davis Technologies | RZ2 |
| Data streamer | Tucker-Davis Technologies | RS4 |
| Diode laser (473 nm) | Omnicon | LuxX-473 |
| DPSS laser (532 nm) | LaserGlow | LRS-0532 |
| Power meter | Thorlabs | S120C |
| Solenoid driver | Eaton-Peabody Lab | https://github.com/EPL-Engineering/epl_valve |
| Lickometer | Eaton-Peabody Lab | https://github.com/EPL-Engineering/epl_lickometer |
| Free-field speaker | Parts Express | 275-010 |
| Titanium headplate | iMaterialise | Custom |

## EXPERIMENTAL MODEL AND SUBJECT DETAILS

### Animal subjects

All procedures were approved by the Massachusetts Eye and Ear Animal Care and Use Committee and followed the guidelines established by the National Institute of Health for the care and use of laboratory animals. Data were collected from 101 male and female mice. To ensure that precocious age-related hearing loss did not interfere with our experiments, we used the PV-Cre line, which naturally retains normal high-frequency hearing through the ages tested here or the Cdh23 line, which features a reverse point mutation in the Cadherin23 gene that causes the accelerated age-related hearing loss in C57 mice. Eight PV-Cre mice were used for the single unit electrophysiology experiments with acute optogenetics. 3 PV-Cre mice contributed to the electrophysiology experiments comparing gamma entrainment across optogenetic and acoustic stimuli. Nine mice from a CDH23 x B6.129S line and 1 PV-Cre mouse were used in the PVN γ stimulation electrophysiology experiments (N = 6/4 control/ γ stimulation). A total of 47 mice contributed to behavioral tasks: 24 to 2AFC post noise-exposure studies (N = 6/12/6, sham/PTS/TTS) and 23 to optogenetic 2AFC behavior (N = 4/14/5, control/ChR2/Arch). Optogenetic activation, inactivation, and control experiments were performed in PV Cre x Ai32,PV Cre mice, and CDH23 x B6.129S mice). Eleven C57BL/6J mice (N = 5/6 sham/PTS) contributed to ABR testing and cortical electrophysiology before and after noise exposure. Cochlear histology data was quantified from 19 mice (N = 7/6/6, sham exposure/SNHL exposure/synaptopathic exposure). Cochlear histopathology data in SNHL mice were originally collected as part of a prior study and reanalyzed here.^55^ Three B6.129S x Cdh23 mice were used for PV immunolabeling experiments. Mice were maintained on a reverse 12 hr light/12 hr dark cycle and were provided with *ad libitum* access to food and water unless they were undergoing behavioral testing, in which case they had restricted access to water in the home cage.

## METHOD DETAILS

### Preparation for behavioral experiments

Mice were anesthetized with isoflurane in oxygen (5% induction, 1.5%–2% maintenance). A homeothermic blanket system was used to maintain body temperature at 36.6° C (FHC). Lidocaine hydrochloride was administered subcutaneously to numb the scalp. The dorsal surface of the scalp was retracted and the underlying periosteum removed. The skull surface was prepped with etchant (C&B metabond) and 70% ethanol. For behavioral subjects not tested with optogenetic manipulations, a titanium head plate (iMaterialise) was then affixed to the dorsal surface of the skull with dental cement (C&B metabond).

For mice undergoing additional optogenetic testing, a small region of the skull overlying the ACtx of each hemisphere was optically cleared. This was accomplished by applying one layer of each in the following order: cyanoacrylate (Krazy Glue), clear dental cement (C&B metabond), and finally clear nail polish (L.A. colors). While the nail polish was drying, we affixed cylindrical opaque mating sleeves for optic fiber assemblies (Doric) atop the cleared skull so that light could efficiently reach the ACtx without implanting a fiber on or in the brain. Once all elements had dried, a headplate was affixed to the skull following the procedure described above. At the conclusion of the headplate attachment for all surgeries, Buprenex (0.5 mg/kg) and meloxicam (0.1 mg/kg) were administered, and the animal was transferred to a warmed recovery chamber.

### Virus injections

For optogenetics experiments, ChR2, Arch, or GFP was expressed in PVNs (ChR2 and Arch) or pan-neuronally (GFP) by making 2-3 small, evenly spaced burr holes in the skull overlying the ACtx of each hemisphere (1.75-2.25 mm rostral to the lambdoid suture) with a 31-gauge needle. Pulled glass micropipettes (Wiretrol II, Drummond) were backfilled with virus solution (ChR2: AAV2/5-Ef1a-DIO -ChR2-EYFP or AAV2/5-S5E2-mCherry, Arch: AAV9-FLEX-Arch-GFP, GFP: AAV2/5-hSyn-EFP), advanced through an individual burr hole, and then 200 nL of virus solution was injected at 1.1 nL/s 0.3 mm below the pial surface using a precision injection system (Nanoject III, Drummond) with a 5 s delay between each injected bolus. At least 10 minutes passed following each injection before the pipette was withdrawn and burr holes were filled with KWIK-SIL (WPI). Injections were performed prior to the skull clearing procedure. For acute electrophysiology experiments, the procedure above was only performed in the right hemisphere.

Mice used in SNHL electrophysiology experiments received additional virus in the left and right auditory cortex (right: AAVrg-pgk-Cre, 250 nL mixed and co-injected with the AAV-S5E2-ChR2 solution; left: AAV2/5-EF1a-DIO-ChrimsonR-mRuby-KV2.1, 350 nL). CDH23 x B6.129S mice used for PV γ stimulation electrophysiology experiments underwent an additional virus injection (300 nL of AV-EF1a-DIO-ChrimsonR-mRuby-KV2.1) into the basal forebrain (A-P: -0.8 mm, M-L: 2.7 mm, D-V: 3.45 mm). In both cases, the Kv2.1 promoter ensured that the red-shifted opsin was expressed only in the somata of Cre-expressing neurons in the contralateral ACtx or ipsilateral basal forebrain with no cell body or axon expression in the right ACtx and hence no influence over any of the data presented here. Findings from experiments arising from these virus manipulations are unrelated to the hypotheses tested here and may be the subject of future publications.

### High-frequency noise exposure

Noise exposure occurred at 10 weeks postnatal and was timed to occur in the morning. For synaptopathic exposure, octave-band noise at 8-16 kHz was presented at 93 dB SPL for 2 hours. For SNHL exposure, octave-band noise at 16-32 kHz was presented for 2 hours at 103/98/70 dB SPL. Exposure stimuli were delivered via a tweeter (Fostex USA) fixed inside a custom-made exposure chamber (51 x 51 x 51 cm). To minimize standing waves, the walls of the acoustic enclosure constructed so that the walls did not form right angles.

Additionally, irregular surface depths and sharp edges were built onto 3 of the 6 walls using stackable ABS plastic blocks (LEGO) to diffuse the high-frequency sound field. Prior to exposure, mice were placed in a wire-mesh chamber (15 x 15 x 10 cm). This chamber was placed at the center of a continuously rotating plate, ensuring mice were exposed to a uniform sound field.

### Auditory brainstem response measurements

Animals were anesthetized with ketamine (120 mg/kg) and xylazine (12 mg/kg), with half the initial ketamine dose given as a booster when required and were placed on a homeothermic heating blanket during testing. Needle electrodes were inserted in the ipsilateral and contralateral pinnae near the tragus and a ground electrode was placed at the base of the tail. ABR stimuli were 5 ms tone pips at 8,12,16 or 32 kHz with a 0.5 ms rise-fall time delivered at 27 Hz in alternating polarity. For each stimulus, responses were averaged over 512 repetitions. Intensity was incremented in 5 dB steps, from 20–100 dB SPL. All noise-exposed mice used for electrophysiology experiments underwent ABR testing. ABR testing was performed on 5/6/6 mice used for behavioral testing before and after SNHL/synaptopathic/sham exposure.

### Cochlear and cortical histology

#### Cochlear histology

Deeply anesthetized animals were transcardially perfused with 4% paraformaldehyde in 0.1 M phosphate buffer, the cochleae were dissected and perfused through the round and oval windows and post-fixed in the same solution for 2 h at room temperature. Cochleae were then decalcified in 0.12 M EDTA for 2 days at room temperature, dissected into half-turns and blocked in 5% normal horse serum and 0.3% Triton X-100 for 1 hour at room temperature. They were immunostained by overnight incubation with a combination of the following primary antibodies: 1) rabbit anti-myosin VIIa (Proteus Biosciences, 1:200); 2) mouse IgG1 anti-CtBP2 (BD Biosciences, 1:200); 3) mouse IgG2a anti-GluA2 (Millipore, 1:2 000) and secondary antibodies AlexaFluor 647 donkey anti-rabbit, 1:200; Alexa Fluor 568 goat anti-mouse IgG1, 1:1 000; Alexa Fluor 488 goat anti-mouse IgG2a, 1:1 000 (Thermo Fisher Scientific) coupled to the blue, red, and green channels, respectively. Immunostained cochlear pieces were measured, and a cochlear frequency map was computed to associate structures to relevant frequency regions using a plug-in to ImageJ. ^61^

#### Cortical histology

Following perfusion, brains were removed and stored in 4% paraformaldehyde for 24 hr before transferring to cryoprotectant (30% sucrose) for 48 hr. Sections (30 µm) were cut using a cryostat (Leica CM3050S), mounted on gelatin-subbed glass slides (BBC Biochemical) and coverslipped. For areal analysis of virus expression, images were obtained with an epifluorescence microscope (Leica). For co-labeling of S5E2-mCherry and PV protein expression (Figure S3), sections were washed in PBS containing 0.1% Triton X-100 (three washes, 5 min each), incubated at room temperature in blocking solution (Super Block) for 6 min at room temperature and then incubated in primary antibody (PV 27 rabbit anti-parvalbumin, Swant,1:1000 dilution) at four degrees C overnight. The next day, slices were washed and incubated in secondary antibody (Alexa Fluor 488 Donkey Anti-Goat IgG, AbCam, 1:200 dilution) for 1.5 hr at room temperature. Fluorescence images were obtained with a confocal microscope (Leica).

### Two-alternative forced-choice task

#### Shaping

Three days after headplate surgery, animals were weighed and placed on a water restriction schedule (1 mL per day). During behavioral training, animals were weighed daily to ensure they remained above 80% of their initial weight and examined for signs of dehydration. Mice were given supplemental water if they received less than 1 mL during a training session or appeared excessively dehydrated. During testing, mice were head-fixed in a dimly lit, single-walled sound-attenuating booth, with their bodies resting in an electrically conductive cradle. Tongue contact on the lickspout closed an electrical circuit that was digitized (at 40 Hz) and encoded to calculate lick timing. Target tones were delivered through an inverted dome tweeter (Parts Express) positioned 10 cm from the left ear. Sound level was calibrated with a ¼” prepolarized microphone (PCB Electronics). Digital and analog signals controlling sound delivery and water reward were controlled by a PXI system with custom software programmed in LabVIEW (National Instruments).

Mice required approximately 3 weeks of behavioral shaping before they could perform the complete 2AFC task. For shaping, mice were first habituated to head-fixation and conditioned with a Pavlovian approach in which 11.3 kHz tones (150ms, 5 ms raised cosine onset-offset ramps) at 40 and 45 dB SPL were presented with a small quantity (3.5 µL) of water on the “soft” spout while 75 and 80 dB SPL tones were paired with water on the “loud” spout. The assignment of loud and soft to the left and right spouts were randomly determined for each mouse. Trials had a variable inter-trial interval (4-10s) selected on each trial from a truncated exponential distribution to produce a flat hazard function. Tone presentation was further delayed to ensure at least 1s had elapsed without lick spout contact. Once the association between sound level and spout was established, mice were advanced to the operant task, in which water delivery was contingent upon licking the correct spout within 1.2s following tone onset. To forestall or mitigate the development of choice bias, a closed-loop training paradigm was intermittently used where mice were repeatedly given the same trial type after an incorrect response until they chose correctly.

Once mice met the criterion of > 90% correct categorization, they were advanced to the full stimulus set, where tone intensity on each trial was randomly selected from a 40 to 80 dB SPL (5 dB step size). At this stage, mice were required to correctly categorize 40-45 dB SPL tones as ‘soft’ and 75-80 dB SPL tones as ‘loud’ by licking the appropriate spouts to receive water, but they were rewarded regardless of their spout choice for 50-70 dB SPL tones. Mice were acclimated to the complete version of the task for 2-3 days, and then behavioral testing began.

#### Testing

Loudness choice functions for a given condition were generated from data collected from 3-4 behavioral sessions. Mice typically performed ∼300 trials per session and 5-7 sessions were performed per week.

#### Loudness testing at a fixed sound level

To understand how sound duration contributes to loudness perception, tone duration (10, 25, 50, 75, 100 ms) was varied at a fixed intensity (60-70 dB SPL) and all choices unconditionally reinforced. To ensure mice continued to perform the loudness classification task, tone duration was only varied on 1/3^rd^ of trials. The remaining trials presented the standard 150 ms tone duration. In other experiments, the intensity of optogenetic PVN stimulation was varied in equally spaced steps up to the maximum intensity (10-15 mW) for a fixed sound intensity (70 dB SPL). As in the duration experiments, PVN stimulation intensity was only varied in one third of trials and responses were unconditionally reinforced.

### Auditory cortex electrophysiology

#### Preparation for acute insertion of high-density probes in awake, head-fixed mice

During the initial head plate attachment surgery, a small burr hole was made over the left occipital cortex and a ground wire was placed. On the day of the recording, the mouse was briefly anesthetized with isoflurane in oxygen (5% induction, 2% maintenance) and small craniotomy (1 x 1 mm) was made on the temporal ridge, 1.5-2.5 mm anterior from the lambdoid suture, to expose the right ACtx. UV-cured composite (Flow-It ALC) was used to create a recording chamber around the craniotomy. The chamber was then filled with optically transparent lubricating ointment (Paralube Vet Ointment) to prevent tissue desiccation and provide recording stability. Isoflurane was then stopped and the mouse was transferred to a dimly lit double walled acoustic chamber where it sat head-fixed in a body cradle. Recordings started at least 30 minutes after the cessation of anesthesia to allow full recovery from isoflurane, as evidenced by normal whisking and grooming behaviors.

Extracellular recordings: A 64-channel silicon probe (H3, Cambridge Neurotech) was slowly advanced (100 µm/s) into ACtx perpendicular to the pial surface until the tip of the electrode was 1.3-1.4mm below the cortical surface, ensuring full coverage of all layers. In experiments characterizing cortical responses after SNHL, two-shank probes were occasionally used (H2, Cambridge Neurotech) to maximize coverage across the deafferented region of the tonotopic map. The brain was allowed to settle for at least 15 minutes before recordings began. On the day of the first recording, multiple penetrations were made to identify the tonotopic reversal which represents the rostral border of A1. For noise exposure experiments, a more detailed tonotopic mapping referenced to a high-resolution image of the cortical surface was performed the day before noise exposure to identify the high-frequency region of A1 for recordings on subsequent days.

Raw neural data was digitized at 32-bit, 24.4 kHz and stored in binary format (PZ5 Neurodigitizer, RZ2 BioAmp Processor, RS4 Data Streamer; Tucker-Davis Technologies). To eliminate artifacts, the common mode signal (channel-averaged neural traces) was subtracted from all channels in the brain. Signals were notch filtered at 60Hz, then band-pass filtered (300-3000 Hz, second order Butterworth filters). To calculate local field potentials, raw signals were first notch filtered at 60 Hz and downsampled to 1 kHz.

Kilosort 2.0 was used to sort spikes into single unit clusters.^95^ Single-unit isolation was based on the presence of both a refractory period within the interspike interval histogram, and an isolation distance (>10) indicating that single-unit clusters were well separated from the surrounding noise.^96^ RS and FS designation was based on the peak to trough delay of the mean spike waveform for each unit, > 0.6 ms (RS) or < 0.5 ms (FS).

#### Acoustic stimuli

Acoustic stimuli were delivered through a freefield tweeter positioned 15-20 cm from the left ear. Output from the speaker was calibrated with a ¼” prepolarized microphone (PCB Electronics). All stimuli were presented in pseudorandom order with 5 ms onset/offset cosine-squared ramps unless otherwise noted.

White noise stimuli (150 ms duration, 0.1-100 kHz,) which varied between 0 to 80 dB SPL in 5 dB steps were used to characterize cortical intensity coding and the effects of acute PVN optogenetic manipulations (20 trials per stimulus, 850 ms inter-stimulus interval (ISI). Population frequency coding was assessed with pure tones ranging from 4-45 kHz in 0.5 octave steps (150 ms duration, 20 to 80 dB in 5 dB SPL steps, 20 trials per stimulus). Sensory-evoked gamma oscillations were measured with a 40 Hz click train (1 μs clicks, 70 dB SPL, 1 s duration, 500 ms ISI). To characterize receptive fields before and after PVN stimulation, tone pips were used (4:45 kHz, 20:5:80 dB SPL, 150 ms duration, 350 ms ISI, 30 trials per stimulus). The full set of tone pip stimuli were presented once before stimulation (baseline), and twice following pairing (+30 and +60 minutes). Acoustic stimulation began immediately following the completion of PVN stimulation. For experiments characterizing sound responses following SNHL, FRAs were first measured to confirm tonotopic position and confirm the presence of a sharp edge in the FRA following high-frequency cochlear deafferentation (4 to 45 kHz in 0.2 octave steps, 10-70 dB SPL in 5 dB steps, 4 trials per stimulus). To characterize intensity coding for spared frequencies, a quarter-octave narrowband noise with an 11.3 kHz center frequency was used (50 ms duration, 0.5 s ISI, 20 trials per stimulus).

### Optogenetic manipulations

#### Optogenetics during loudness classification behavioral task and PVN stimulation

Trials with sound and light stimulation (33%) were randomly interleaved with sound-only trials. Light was delivered to the brain via an optic fiber/ferrule assembly (0.2 mm diameter, 0.22 NA Doric) coupled to a 473 nm diode laser for ChR2 activation (Omnicron LuxX) or a 532 nm laser for Arch activation (LaserGlow). The light path was split to provide a light source for each hemisphere separately. Laser power was individually calibrated for every fiber to produce 10 mW at the fiber tip. Light transmittance through the skull was measured *ex vivo* in a subset of mice and found to be approximately 20% of the input power. The voltage command signal to the laser specified a 25 ms pulse widths over a 0.6 s period at 50% duty cycle (i.e., 20Hz) shaped with a tapering amplitude envelope (0.1s) to reduce any rebound excitation. A black mating sleeve (Doric) was attached to each laser patch cable, which fit snugly into the skull-mounted plastic casings. Laser onset preceded tone onset by 100 ms. For PVN stimulation (γ or control) experiments, mice were placed in the behavioral chamber with the lick spouts removed and subjected to 16 minutes of bilateral 473 nm light stimulation (γ: 10 mW, 40 Hz stimulation rate,1 ms pulse width, 1 s trial duration, 1000 trials; control: 10 mW, 50 ms pulse width, 1 s ISI, 1000 trials).

#### Optogenetic during ACtx electrophysiology

For electrophysiology experiments, a laser-coupled optic fiber was advanced to 1-2 mm above the surface of the exposed ACtx. For experiments interleaving sound with versus without laser stimulation on neural coding of sound level, stimulation parameters and hardware were identical to behavioral manipulations, except that 30 mW was required for Arch experiments, consistent with previous literature.^40^ To study PVN gamma entrainment, 473 nm laser stimulation was either constant or pulsed at 40Hz for a 1s period, using the presentation parameters described above. To study PVN-mediated suppression of RS unit activity, a 0.5s period of variable power with a 0.1s offset taper was pseudorandomly presented every 1.2s until each had been presented 20 times.

## QUANTIFICATIONS AND STATISTICAL ANALYSIS

### Electrophysiology analysis

#### Rate-level functions

A unit was determined to be sound-responsive by performing a paired t-test between trial-by-trial spiking 50 ms before and 8 to 58 ms post stimulus onset across all acoustic stimuli. Rate level functions were computed for all sound-responsive units by subtracting the mean spiking 8 to 30 ms post-stimulus onset from the mean spiking during a 0.5 s baseline period ending 0.1s prior to stimulus onset (to avoid including optogenetic changes in spiking in the baseline rate). For visualization and clustering, rate-level functions were normalized to their absolute maximum. K-means clustering was performed on all sound-responsive units in Matlab using the “kmeans” function (maximum iterations, 1000; replicates, 10; distance metric, correlation). For the gain analysis following SNHL, spontaneous activity collected during a 200ms pre-stimulus baseline was subtracted from spiking occurring during a 25ms period beginning at the first post-stimulus spike. The area under the rate-level function above response threshold was computed using the “trapz” function in Matlab.

#### Sound level decoding

The input to population-based classifiers were the raw spike rates from 8 to 38 ms post-stimulus onset for a randomly drawn set of 250 RS units, chosen regardless of sound responsiveness. In the population rate model, the summed population spike rate for each trial was used as the input to a leave-one-out classifier. For a given sound level or frequency, the population spike rate on the held-out trial was compared to the mean population rates across all stimuli in the training set and the classified stimulus was defined as the most similar population spike rate. For the higher-dimensional classifier, principal components analysis was first performed on the unit by trial response matrix, and the first 5 principal components were used to decompose response variations across trials, sound levels, and individual units among the 250 RS unit ensembles. The leave-one-out procedure was similar to the population rate model, but the classified stimulus was assigned based on minimizing the Euclidean distance between the 5-D trial response vector and the average response vectors for each stimulus. Classification was repeated until each trial was used as the held-out test trial. Overall classification was averaged across 1000 random draws of RS units. Chance performance was assessed for each model by performing classification on with shuffled stimulus labels (50 stimulus shuffles, each with 1000 random draws of RS units). To assess the effects of acute PVN activation or inactivation on intensity classification, individual laser-on trials were decoded using laser-off training data.

#### LFP and single unit gamma rhythm analysis

For LFP analysis, the noise-evoked pattern of sinks and sources in the current source density, the second spatial derivative of the LFP) were used to identify electrode channels within versus outside L1-6 of A1. The early Layer 4/5 sink which appears around 8-10 ms post stimulus onset was manually identified and the corresponding channel depth was set as 496 μm below the pial surface. Only channels below the pial surface or above the white matter (<1 mm) were considered within ACtx. To quantify gamma entrainment in the trial-averaged LFP, the short-time Fourier transform was computed for each channel within A1 and then averaged (80 ms Hanning window, 79 ms overlap). Entrainment was defined as the relative power in the 35-45 Hz stimulus-evoked gamma band compared to other frequencies (<30 Hz and 55-70 Hz) which did not include harmonics of the stimulus frequency. Power in each frequency band was referenced to its own pre-stimulus baseline (250 ms). For spiking entrainment, the short-time Fourier transform was computed on the average spike rate across the recorded population of single units for each penetration, as individual units could have near-zero average spike rates. Relative gamma entrainment for spiking was calculated using the same pre-stimulus baseline and frequency ranges as for the LFP analysis.

To analyze changes in spontaneous LFP power, epochs without sensory or optogenetic stimuli (500 ms duration, 120 trials for PVN stimulation experiments, 700 ms duration, 50 trials for SNHL experiments) were filtered between 0.1 and 300 Hz using a second-order Butterworth filter. The fast-Fourier transform was performed to extract the on the single-trial LFP power from each channel following tapering using a Hamming window equivalent to the trial duration. Channels included in the analysis were spaced every 200 μm across the depth of auditory cortex to avoid artificial sample size inflation by non-independent samples and all other channels within A1 were excluded. Channels below the white matter were labeled as non-ACtx and again sampled at 200 μm resolution, while channels above the pia were excluded. Delta and gamma bands was defined as 2-4 and 30-80 Hz, respectively.

#### Short-term plasticity of sound-evoked spiking activity following bouts of PVN stimulation

FS units were split at the 50^th^ percentile into two groups (FS_entrained_ vs. FS_non-entrained_) based on their evoked spiking rate during the PVN stimulation period, referenced to the pre-stimulation baseline. Sound-evoked spiking for each frequency-level combination was computed as the mean of activity 8 to 50 ms post-stimulus onset referenced to the mean of activity in the 100 ms pre-stimulus. Only units that were sound-responsive were included for unit-level analyses. A unit was classified as sound-responsive using a paired t-test on the trial-by-trial spiking activity 100 ms pre-stimulus compared to 8 to 50 ms post-stimulus. Best frequency was defined as the frequency that elicited the maximal spiking across all sound levels. Rate-level functions were taken as the mean evoked rate at BF ± 0.5 octaves for each sound level. Evoked activity was averaged across a 50-80 dB range and change in sound response was calculated as the fold change (pre/post). All RS units were used for the population rate classifier, however only 100 units were used in each population draw due to a lower sample size. Procedures for classifier training and testing were identical; here, intensity classification relied on single trials from the post-stimulation period referenced to pre-stimulation data.

#### Assessing the strength of PVN-mediated inhibition

To estimate the strength of PVN-mediated feedforward inhibition, an asymmetry index was calculated in RS units as the mean spike rate [post-pre] / [post+pre], where the pre-stimulus period was 500 ms and the post-stimulus period was the first 200ms of the 500ms continuous laser period. To prevent ceiling or floor effects, only laser powers where the baseline asymmetry index for each was less than +/-0.75 were considered for further analysis. For the SNHL experiments, the asymmetry index was calculated based on the mean spike rates during the complete 500 ms stimulation period as [XmW – 0mW]/ [XmW + 0mW] where X refers to all laser powers above the 0mW control . To calculate the fraction of FS or RS units activated by laser stimulation for S5E2 vs. Cre-dependent ChR2 expression, a unit was considered activated if 10 mW laser stimulation increased its mean firing rate by >5 standard deviations above its pre-stimulus baseline.

#### ABR

ABR threshold was defined as the lowest stimulus level at which a repeatable waveform could be identified. ABR wave amplitude was defined as the peak amplitude – the mean absolute value during a 0.1s period prior to stimulus onset.

### Behavioral analysis

Psychometric functions were fit to raw left/right choice data using binary logistic regression. To get the fits across time or across mice, raw data was first concatenated. The loudness transition point was defined as the intensity where the psychometric function crossed a 50% loudness choice probability. Reaction time was defined as the first lick following stimulus presentation. Reaction times were averaged across all intensities and expressed relative to baseline.

### Histological analysis

#### Virus expression

Boundaries of virus expression were calculated from serial coronal sections registered to the Allen Brain Atlas and identified with AP-Histology toolbox. To assess the colocalization of PV and S5E2-ChR2, a region of interest was drawn around the area of viral expression in ACtx and cells within the area which expressed S5E2-ChR2 and PV, PV alone, or S5E2-ChR2 alone were manually counted by two independent observers.

#### Inner hair cell synaptic innervation

Confocal z-stacks at 8.4, 16, 22.6, 32 and 45 kHz frequency locations were collected using a Leica TCS SP8 microscope using a glycerol-immersion objective (63x, N.A. 1.3) and 2.38x digital zoom to visualize inner hair cells (IHCs) and synaptic structures. Two adjacent stacks were obtained at each target frequency spanning the cuticular plate to the synaptic pole of ∼10 hair cells (in 0.33 µm z-steps).

Images were loaded into an image-processing software platform (Amira; ThermoFisher Scientific), where IHCs were quantified based on their Myosin VIIa-stained cell bodies and CtBP2-stained nuclei. Presynaptic ribbons and postsynaptic glutamate receptor patches were counted using 3D representations of each confocal z-stack.

Juxtaposed ribbons and receptor puncta constitute a synapse, and these synaptic associations were determined by calculating and displaying the x–y projection of the voxel space within 1 µm of each ribbon’s center . Mean counts for the sham group were used to normalize all IHC ribbon synapse data.

### Statistical analysis

All statistical analyses were performed in MATLAB 2021b (Mathworks). Details of all statistical testing are provided in the figure legends. Post hoc pairwise comparisons were corrected for multiple comparisons using the Holm-Bonferroni correction. Unless otherwise stated, error shown is the standard error of the mean (SEM).

## Supplemental Information

**Figure S1.**
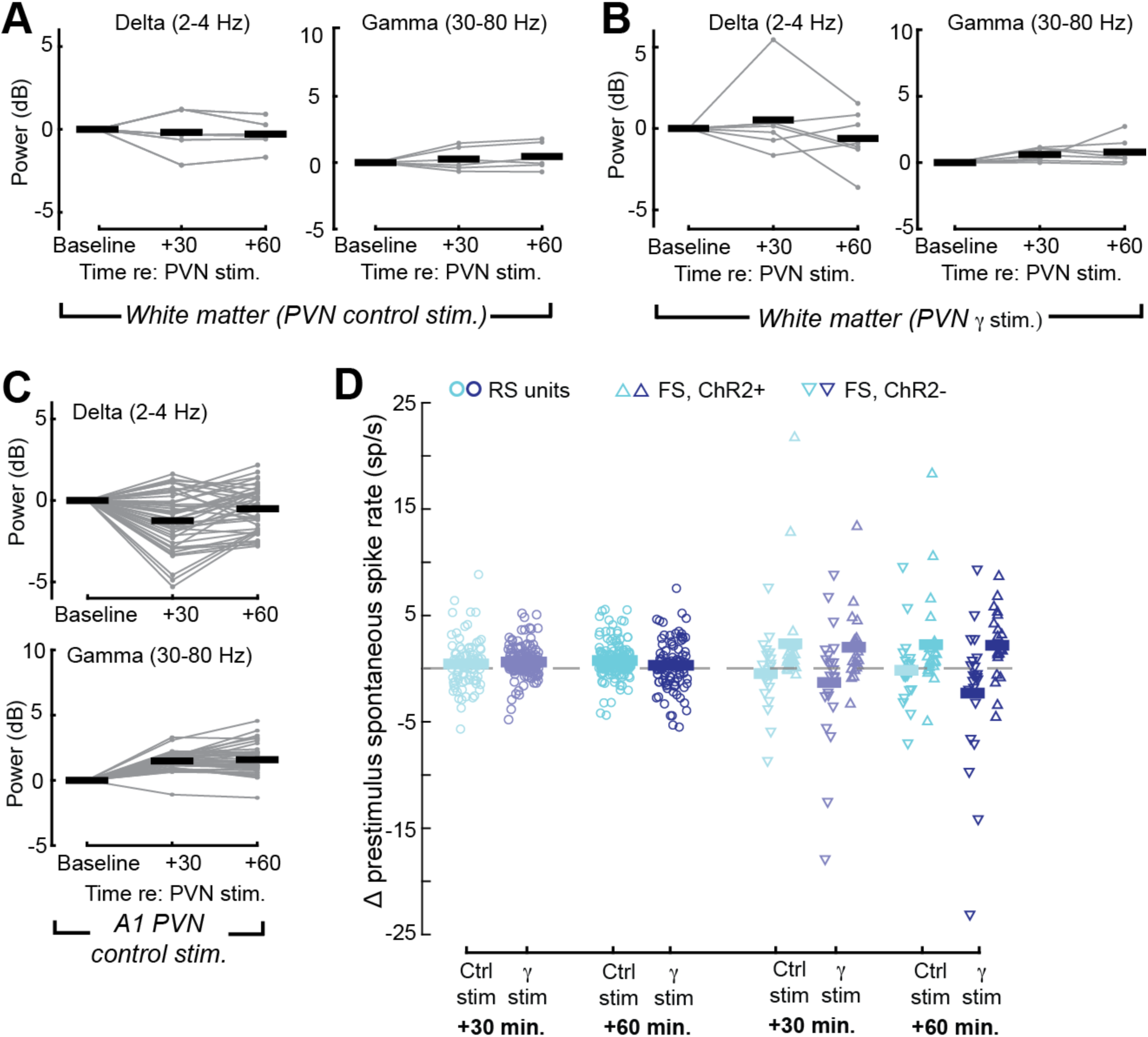
PVN γ and control stimulation persistently change spontaneous LFP power but not spike rate. (A) Change in gamma and delta power for electrode sites outside ACtx after 16-minute bout of PVN control stimulation (N = 6 mice, n = 7 sites). Thin lines depict individual electrode sites. Gamma and delta power was unchanged at 60 minutes (t-test against a population mean of zero, delta: t = -0.98, p = 0.57; gamma: t = 2.07, p = 0.08). (B) Change in gamma and delta power for electrode sites outside ACtx after PVN γ stimulation (N = 4 mice, n = 6 sites). Gamma and delta power was unchanged at 60 minutes (t-test against a population mean of zero, delta: t = -0.34, p = 0.75; gamma: t = 0.76, p = 0.48). (C) Change in gamma and delta power for electrode sites in ACtx after control PVN control stimulation (N = 6 mice, n = 42 sites). Gamma power was increased (t-test against a population mean of zero: t = 13.82, p = 5 x 10^-17^) and delta power was decreased (t = -4.54, p = 4.78 x 10^-5^). (D) Change in spontaneous rate firing rate of sound responsive single units following PVN stimulation. Spontaneous rates increased on average (mixed-design ANOVA, main effect for time, F = 6.24, p = 0.013, main effect for cell-type, F = 17.59, p = 5.42 x 10^-8^, Cell type x Time x Stimulation type, F = 1.73, p = 0.178), but changes were similar between stimulation groups (unpaired t-test comparing change in spontaneous rates at 60 minutes: t = 1.87, p = 0.06).

**Figure S2.**
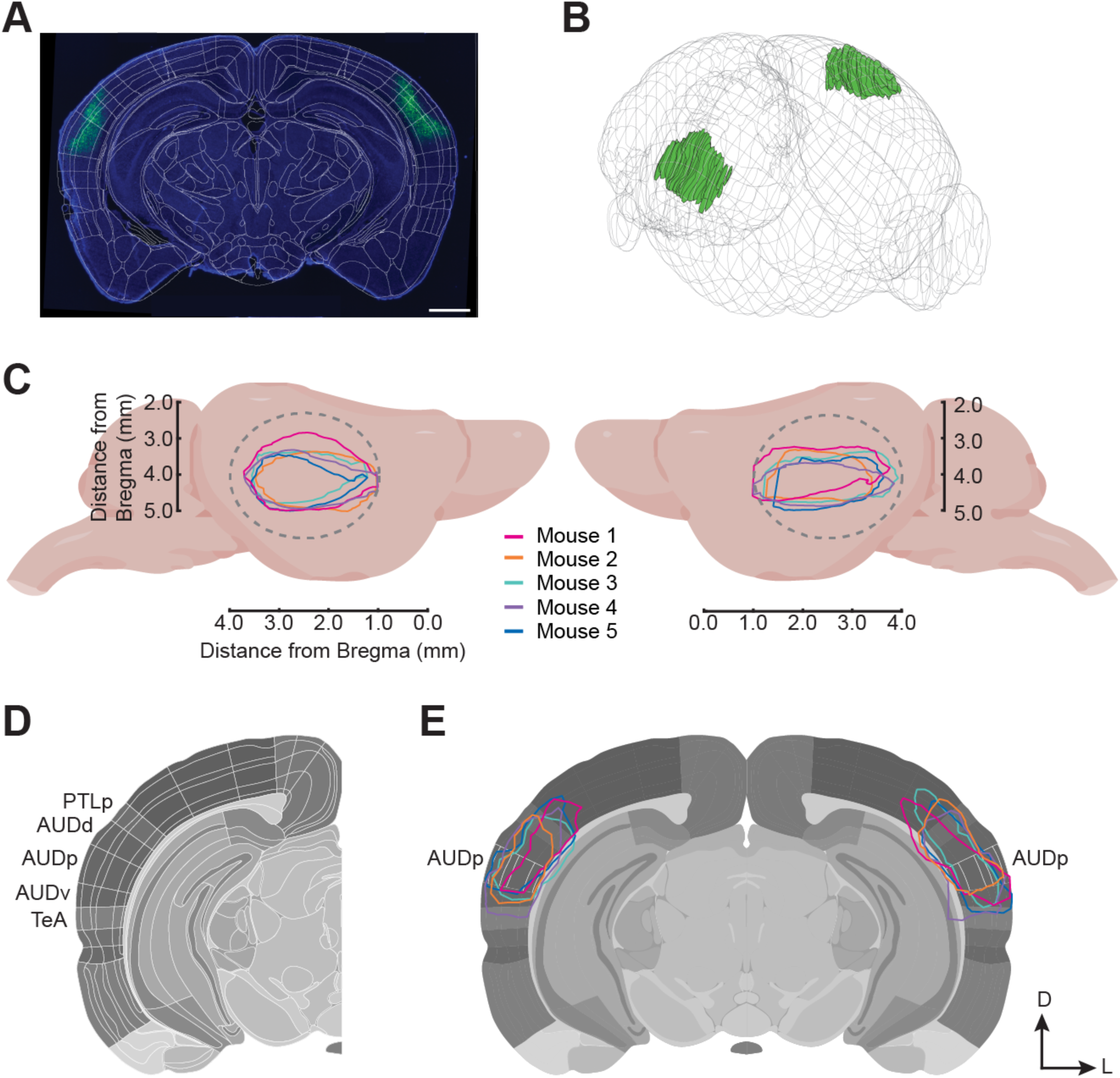
Reconstruction of virus-mediated opsin expression in cortical PVNs. Related to Figure 5-8. A) Example coronal slice (30 µm) registered to the Allen Brain common coordinate framework; DAPI is shown in blue, AAV5-DIO-ChR2-eYFP is shown in green. Scale bar indicates 1 mm. B) Viral expression was manually annotated in each coronal slice. Bilateral expression patterns are shown in a wireframe brain by plotting the labeled area (green) in each coronal slice separately. C) The rostral-caudal and medio-lateral spread of virus expression is shown for each mouse (N = 5) that underwent noise exposure and 40 Hz PVN stimulation. Mouse brain cartoon is not to scale. Gray dotted line indicates the approximate coverage of the clear skull preparation. D) Delineation of cortical layers and fields in the Allen Brain atlas. PTLp = posterior parietal association area; AUD = auditory cortex; p/d/v = primary/dorsal/ventral; TeA = temporal association area. E) The coronal section with the largest spread of virus expression across five mice demonstrates transduction of all layers in all fields of the auditory cortex (3.0 mm ± 250 µm posterior to bregma). Scale bar is 1 mm. White outline denotes AUDp. D and L = dorsal and lateral.

**Figure S3.**
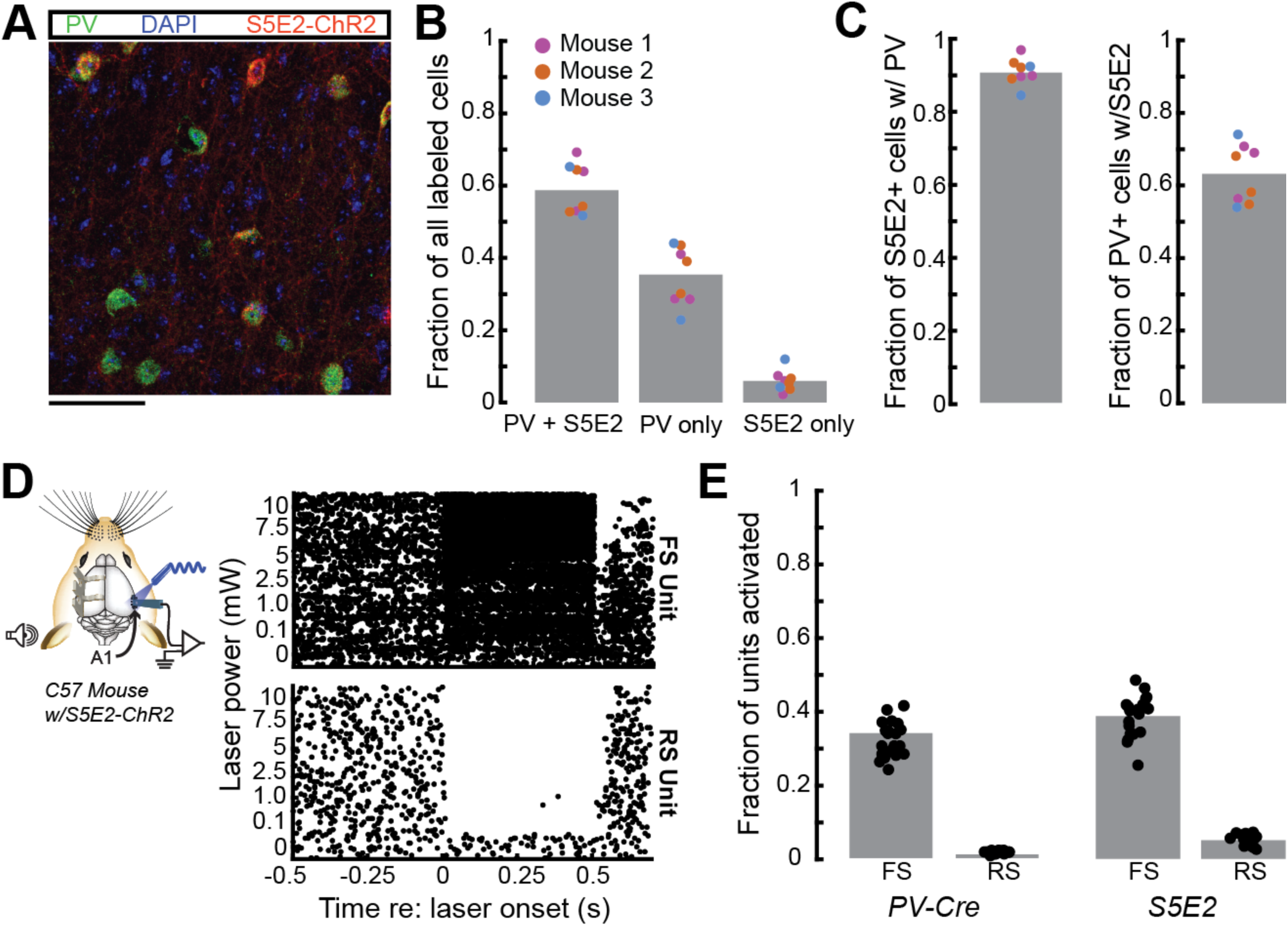
The S5E2 enhancer allows for PVN targeting. Related to Figure 5-7. A) Representative micrograph of ACtx from a WT mouse three weeks post AAV2/5-S5E2-ChR2-mCherry injection (red). Immunohistochemistry was performed to identify cells that express PV and DAPI was used to visualize cell nuclei. Scale bar is 50 µm. B) S5E2 and PV expression was quantified across 3 mice (8 slices, 826 cells). Data points show individual slices and are colored by mouse. 58.72 % of cells were labeled for both S5E2-mCherry and PV, 35.35% expressed PV alone, and 5.93% expressed S5E2-mCherry alone. C) The specificity of S5E2-mediated expression for PV cells was assessed by quantifying the fraction of S5E2-positive cells that express PV (91.00 ± 1.29%). The sensitivity of S5E2-mediated viral expression across the PV+ population was quantified as the fraction of PV cells that express S5E2 (63.16 ± 2.87%). D) Alongside histological quantification, S5E2 expression was also quantified using extracellular recordings where it is possible to identify putative-PV+ units based on their spike waveform. Right, representative FS and RS unit rasters in response to 465 nm laser stimulation of varying power (500 ms, continuous pulse) in a WT mouse expressing S5E2-ChR2. E) The specificity of S5E2-based viral expression was calculated as the fraction of FS and RS units which were significantly activated (laser response > 5 STDs re: baseline) by 10 mW static blue laser stimulation for WT mice with S5E2-ChR2 expression (N = 9, N = 578 units) and PV-Cre mice with Flex-ChR2 (N = 3, N = 376 units). The fraction of FS and RS units activated was comparable across the viral approaches, though there was a slight increase in both FS and RS activation using the S5E2 approach (PV-Cre: FS, 34.48 ± 4.78%, RS, 1.62 ± 0.35%; S5E2: FS, 39.00 ± 5.58%, RS, 5.01 ± 1.47%). Individual data points were obtained by subsampling populations of 250 units from the total populations 20 times as FS unit numbers varied substantially between penetrations, with as few as zero FS units/penetration.

**Figure S4.**
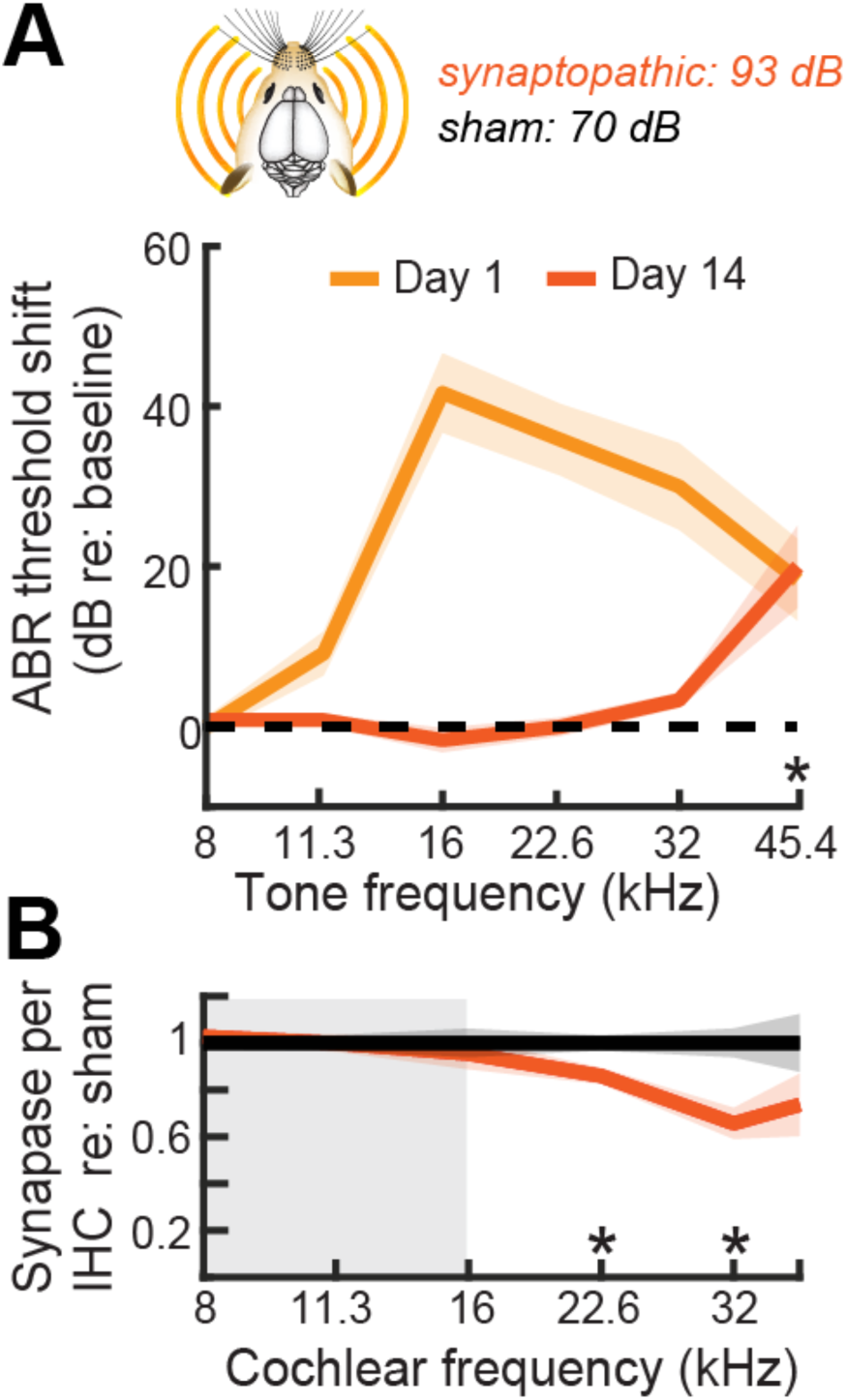
Moderate intensity noise exposure causes a permanent loss of afferent synapses in the cochlear base but a temporary shift in ABR thresholds. Related to Figure 6. (A) Mice were exposed to a high-frequency octave band noise for 2 hours at either 93 or 70 dB. The moderate noise level produced a transient shift in ABR wave 1 thresholds that returned to near baseline for all but 45 kHz at two weeks post exposure (2-way repeated measures ANOVA: Frequency term [F = 10.53, p = 1.6 x 10^-5^] and Time term [F = 47.2, p = 3.2 x 10^-10^]). Asterisks denote significant differences between baseline and day 14 with post-hoc comparisons (p < 0.05). (B) Two weeks after exposure to 93dB noise, loss of synapses between IHCs and auditory nerve fibers was observed in the high frequency region of the cochlea compared to sham exposed ears (2-way repeated measures ANOVA: Frequency x Group interaction, F = 4.2, p = 0.003). Asterisks denote significant differences between sham and synaptopathic noise exposures (p < 0.05).

